# SpatialDE2: Fast and localized variance component analysis of spatial transcriptomics

**DOI:** 10.1101/2021.10.27.466045

**Authors:** Ilia Kats, Roser Vento-Tormo, Oliver Stegle

## Abstract

Spatial transcriptomics is now a mature technology, allowing to assay gene expression changes in the histological context of complex tissues. A canonical analysis workflow starts with the identification of tissue zones that share similar expression profiles, followed by the detection of highly variable or spatially variable genes. Rapid increases in the scale and complexity of spatial transcriptomic datasets demand that these analysis steps are conducted in a consistent and integrated manner, a requirement that is not met by current methods. To address this, we here present SpatialDE2, which unifies the mapping of tissue zones and spatial variable gene detection as integrated software framework, while at the same time advancing current algorithms for both of these steps. Formulated in a Bayesian framework, the model accounts for the Poisson count noise, while simultaneously offering superior computational speed compared to previous methods. We validate SpatialDE2 using simulated data and illustrate its utility in the context of two real-world applications to the spatial transcriptomics profiles of the mouse brain and human endometrium.

## Introduction

Spatial transcriptomics technologies have gained considerable attention, as they enable the exploration of cellular identity and function in a cell’s local environment. Driven by rapid technological evolution, they now allow for the study of hundreds or thousands of genes in parallel. Technologies based on multiplex imaging, such as SeqFISH+ or MERFISH (Eng *et al*, 2019; Xia *et al*, 2019), allow for assaying hundreds of molecular parameters at the same time, providing subcellular resolution. Methods based on RNA sequencing (RNAseq) use spatially barcoded primers and allow a resolution of a handful of cells and approaching single-cell resolution. These methods include Spatial Transcriptomics (Ståhl *et al*, 2016), HDST (Vickovic *et al*, 2019), and Slide-seq (Stickels *et al*, 2021), with a resolution of a handful of cells and approaching single-cell resolution. In particular, 10x Visium, an improved version of Spatial Transcriptomics, is seeing wide adoption in the community due to its commercial availability and ease of use.

While initial studies were focused on proof-of-concept applications in small relatively homogeneous tissue sections, these technologies are increasingly applied to complex tissues or organs with distinct structure and regions, including the human brain, endometrium, and the tumor microenvironment (Kleshchevnikov *et al*, 2020; Garcia-Alonso *et al*, 2021; Baccin *et al*, 2020; Carlberg *et al*, 2019; Thrane *et al*, 2018; Foster *et al*, 2021; Hudson & Sudmeier, 2021; Chen *et al*, 2021). While in principle these data permit addressing a range of distinct questions, a canonical starting point of the spatial omics analysis workflows is the identification of spatially variable genes (Maynard *et al*, 2021; Wang *et al*, 2020b). This step yields genes that are most relevant for downstream analyses, such as the definition of local niches in the tissue that support cellular differentiation and function, but knowledge about spatial variability in its own right can often provide biological insights, e.g. into cancer (Thrane *et al*, 2018). However, existing methods for spatially variable gene detection typically consider the whole field of view within spatial transcriptomics dataset. As the field of view of spatial transcriptomics is growing and the methods are applied to increasingly complex tissues consisting of regions with different cell type compositions, naively applied spatially variable gene detection can no longer yield relevant insights, since the set of identified spatially variable genes primarily contains cell type markers that are not inherently spatially variable (Cable *et al*, 2021). Thus, spatial variance analysis needs to be coupled with suitable computational methods for the identification of tissue regions.

Both spatially variable gene detection and identification of tissue regions have received attention in the field. For example, SpatialDE1 (Svensson *et al*, 2018) was one of the first computational solutions to detect spatially variable genes, which more recently has been refined by the SPARK model (Sun *et al*, 2020). Both models are based on non-parametric Gaussian Process (GP) regression, with SPARK additionally offering a count-based likelihood and a more powerful statistical test. SVCA (Arnol *et al*, 2019), also based on GP regression, and scHOT (Ghazanfar *et al*, 2020), based on distance-weighted nonparametric regression, extend the principles of spatial variable gene detection to provide a more fine-grained decomposition of spatial gene expression variation by accounting for interactions between neighboring cells or voxels. All of these methods have in common that they do not scale to large data sets however, in part due to CPU-bound implementations that cannot take advantage of modern highly parallelized GPU architectures, and in part due to inefficient algorithms. Similarly, a range of clustering methods exists, but they are not well suited for identifying tissue regions in spatial gene expression data. ScanPy (Wolf *et al*, 2018) and Seurat (Stuart *et al*, 2019), two widely used frameworks for scRNA-seq analysis, recommend Leiden clustering to identify tissue regions from spatial omics, an algorithm that is unaware of spatial relationships. Giotto (Dries *et al*, 2021) proposes a method based on hidden markov random fields for this task, thereby imposing a spatial smoothness constraint. However, this model requires the user to pre-specify the number of tissue regions and it employs a Gaussian likelihood model that is suboptimal for count data. Finally, we note that there is a lack of integrated software and workflows to be able to combine spatial variable gene selection and the identification of tissue regions.

Here, we present SpatialDE2, a flexible framework for modeling spatial transcriptomics data. SpatialDE2 implements two major modules, which together provide for an end-to-end workflow for analyzing spatial transcriptomics data: a tissue region segmentation module and a module for detecting spatially variable genes. The tissue region segmentation is fast and provides improved usability compared to previous methods. In particular, the module is capable of automatically determining the number of tissue regions while employing an appropriate count-based likelihood. The module for the detection of spatially variable genes extends previous methods such as SpatialDE and SVCA by providing technical innovations and computational speedups. Calculations of SpatialDE2 can be accelerated using GPUs, opening up the possibility to analyse larger datasets. We validate SpatialDE2’s segmentation and spatial variable gene detection modules using simulated data and application to two real-world datasets, containing 10x Visium data of the mouse brain and human endometrium. In these applications, we demonstrate improved speed and robustness of both modules compared to previous approaches. SpatialDE2 provides an integrated solution for assessing spatial expression heterogeneity within tissue regions, thus providing a principled strategy for dealing with complex tissues and large samples.

## Results

SpatialDE2 implements an end-to-end workflow for the characterization of sub-regional spatial heterogeneity, by implementing two seamlessly integrated analysis modules: a *tissue region segmentation* module, and a module for *spatially variable gene detection*. (**Fig. 1A**). Both modules directly model raw mRNA counts, as obtained from de-multiplexed spatial transcriptomics workflows, or imaging technologies as input. Optionally, SpatialDE2 can also operate on cell count estimates obtained from an additional deconvolution step (Kleshchevnikov *et al*, 2020; Biancalani *et al*, 2020; Cable *et al*, 2021; Lopez *et al*, 2021; Andersson *et al*, 2020; Elosua-Bayes *et al*, 2021).

**Figure 1.**
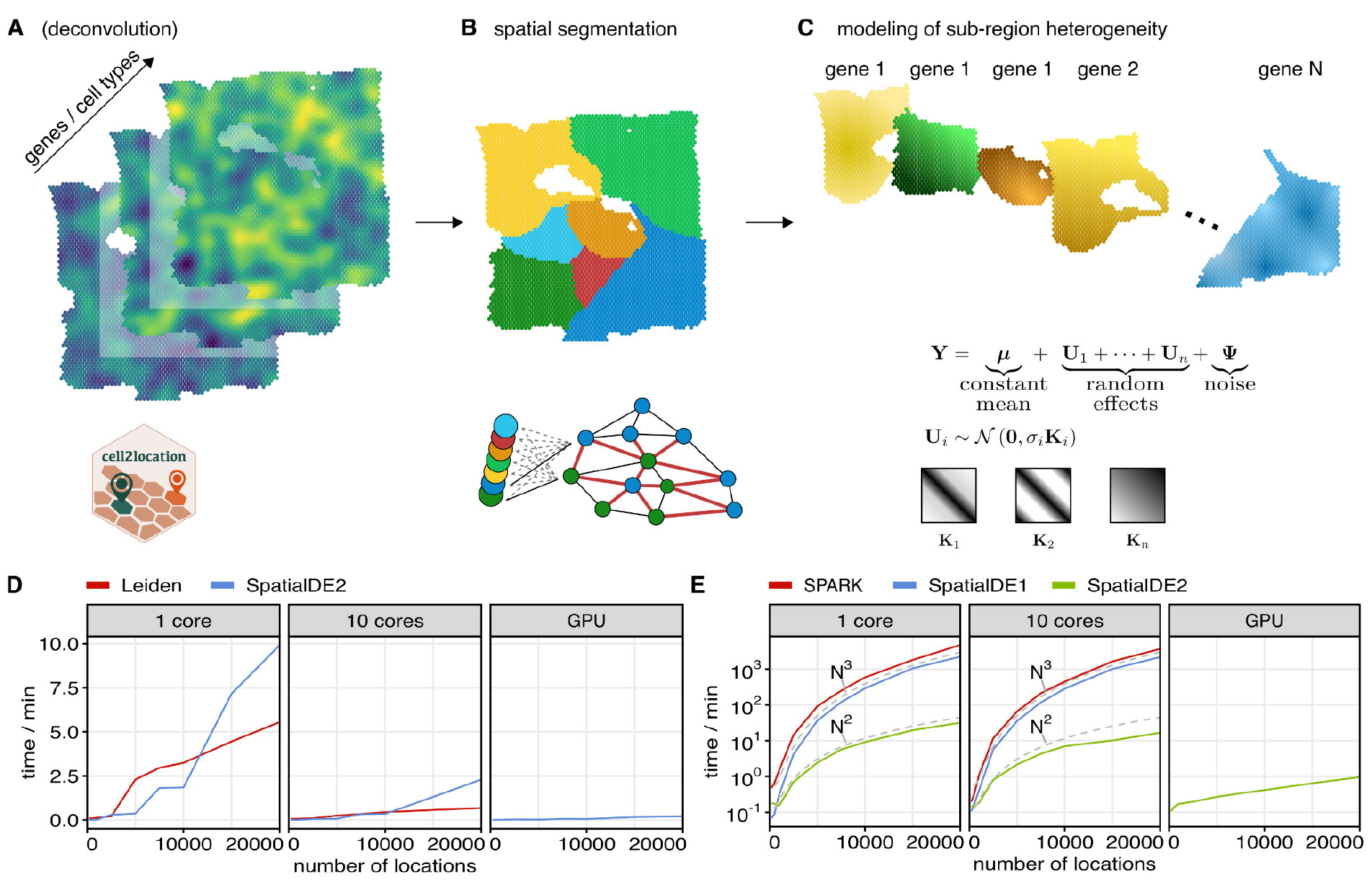
Overview of SpatialDE2 and its computational complexity. (A) Top: SpatialDE2 accepts input spatial transcriptome profiles from a tissue sample, either in the form of a raw gene count matrix, or using cell type counts as provided from a computational deconvolution step (e.g. cell2location). Bottom: input data processing workflow. (B) SpatialDE2 segments the input tissue into connected and transcriptionally similar regions. Top: Schematic output of the segmentation with colours denoting identified tissue regions. Bottom: implementation of the spatial segmentation based on a Poisson Hidden Markov Random field to encode the assumption of spatial smoothness. Nodes correspond to locations with colour denoting the region label. The number of regions is determined by the model. (C) SpatialDE2 models spatial variance components of individual genes within tissue regions. Top: Schematic for the identification of spatial variance components of individual genes in specific tissue regions. Bottom: SpatialDE2 models expression variation within tissue regions by partitioning gene expression variation into one or multiple functional components (U_1_,..,U_n_). Each component is characterized by a covariance matrix that is parametrized by spatial or non-spatial covariates. The special case of spatially variable gene selection corresponds to a functional component parametrized by distance between locations. (D) Run time of SpatialDE2’s tissue region segmentation module and a Leiden clustering workflow for semi-synthetic dataset of increasing size and when using alternative compute environments. Clustering/segmentation was based on 2,000 genes. Leiden denotes the scanpy workflow (Methods). Only SpatialDE2 supports GPU computations. (E) Run time of SpatialDE2’s spatially variable gene selection module versus alternative methods. Considered is a dataset consisting of 200 genes for increasing numbers of locations and alternative computing environments. Only SpatialDE2 supports GPU computations.

Briefly, the spatial tissue region segmentation module is based on a Bayesian hidden markov random field, which segments tissues into distinct histological regions while explicitly accounting for spatial smoothness between neighbouring locations (**Fig. 1B, Methods**). The key advantages of this approach compared to the currently most widely used workflows based on Leiden clustering (Wolf *et al*, 2018; Palla; Stuart *et al*, 2019; Butler; Garcia-Alonso *et al*, 2021) are twofold. First, unlike previous methods SpatialDE2 explicitly accounts for the spatial information during the segmentation step. Second, SpatialDE2 provides a coherent model for tissue segmentation implemented using a principled Bayesian approach, which requires a single user-defined parameter that encodes *a priori* assumptions on spatial smoothness (**Methods**). In contrast, the results from a Leiden clustering workflow are affected in non-intuitive ways by the interplay of manually defined processing steps, each of which is determined by multiple parameters. A common workflow that is referred to hereafter as the Leiden clustering workflow consists of raw data normalization, dimensionality reduction, k-nearest neighbor search, and the application of the Leiden clustering algorithm itself (**Methods**).

The spatially variable gene detection module models variance components of individual genes within identified regions using an appropriate count-based likelihood. While SpatialDE2 builds on Gaussian process regression, which is also used by methods such as SpatialDE (Svensson *et al*, 2018), SVCA (Arnol *et al*, 2019), and SPARK (Sun *et al*, 2020), the model generalizes these approaches by providing novel features, improved scalability, supporting multiple variance components and GPU computations (**Methods, Table S1**). At the core, the model decomposes the expression variation into distinct structured components and a noise term that models random variability (**Fig. 1C, Methods**). Depending on the design of one or multiple covariance matrices, different tests can be formalized that either quantify spatially variable genes, or identify genes that are regulated by cell-cell interactions. Detection of spatially variable genes is implemented as a special case that permits further optimizations (**Methods**).

Finally, we note that SpatialDE2 provides downstream analysis tools to aid interpretation. This includes an updated version of automated expression histology (AEH) (Svensson *et al*, 2018) to identify groups of genes that covary in space. AEH models expression of a particular gene as coming from one of a defined number of smooth spatial patterns and attempts to estimate both the patterns and the assignment of genes to patterns using a Bayesian framework. The number of spatial patterns has to be specified by the user in SpatialDE and can be determined automatically by SpatialDE2.

SpatialDE2 is implemented in Tensorflow (Abadi *et al*, 2016) and supports GPU computations for efficient processing of larger data sets. Run time on a CPU is similar or faster than alternative methods, whereas GPU acceleration not supported by previous methodology enables orders of magnitude faster computations (**Fig 1 D, E, Supplementary Figure S1C**).

## Model validation using simulated data

First, we used simulated data under the null hypothesis of no spatially variable gene expression (**Methods**) to confirm the statistical calibration of the spatially variable gene detection module (**Supplementary Figure S1A**). Briefly, similar to SpatialDE and SPARK, SpatialDE2 increases the sensitivity of its spatially variable gene detection by testing multiple kernel matrices for each gene. SpatialDE2 implements two strategies to estimate statistical significance: Each kernel matrix is tested separately and the p-values are combined using the Cauchy combination (Liu & Xie, 2020), or all kernel matrices are tested simultaneously using an omnibus test (**Methods**). The latter option is faster, since only a single test is conducted, however this strategy results in marginally reduced power (**Supplementary Figure S1B)**. We confirmed that both strategies yielded calibrated results (**Supplementary Figure S1A**). The omnibus test was used for all analyses in this paper.

Next, we simulated true spatially variable genes with variable length scales, adapting empirical parameters from a 10x Visium mouse brain data set (**Methods**). We assessed the sensitivity (statistical power) of SpatialDE2, SpatialDE, and SPARK to detect true spatially variable genes. SpatialDE had the lowest sensitivity, whereas SpatialDE2 and SPARK yielded comparable results (**Supplementary Figure S1B**). By default, SPARK enforces positive definiteness of the kernel by performing an Eigen decomposition of the kernel matrix and setting negative eigenvalues to zero. Since its default periodic kernel is not positive definite, this results in poorly defined and uninterpretable kernels. We therefore also included SPARK without eigenvalue clipping in our benchmark, observing consistent results.

## Tissue region segmentation recovers known histological features along a continuous resolution gradient

Next, we considered 10x Visium data from the mouse brain (Kleshchevnikov *et al*, 2020) to assess SpatialDE2 and alternative methods for tissue segmentation. The mouse brain is well suited for such benchmarking purposes, as it features well-defined and well-annotated distinct regions. Using default settings on spatial smoothness, SpatialDE2 correctly resolved major brain regions (**Fig. 2A, B**), and in particular separated the hippocampal pyramidal/granule cells from the surrounding hippocampal formation (region 0) and identified the lateral ventricle (region 3). For comparison, we also considered a Leiden clustering workflow (**Methods**) applied to the same input data. The segmentation solutions as obtained from Leiden clustering failed to resolve some of the expected brain regions, where in particular both the hippocampal pyramidal/granule cells and the lateral ventricle were not identified (**Fig. 2A**).

**Figure 2.**
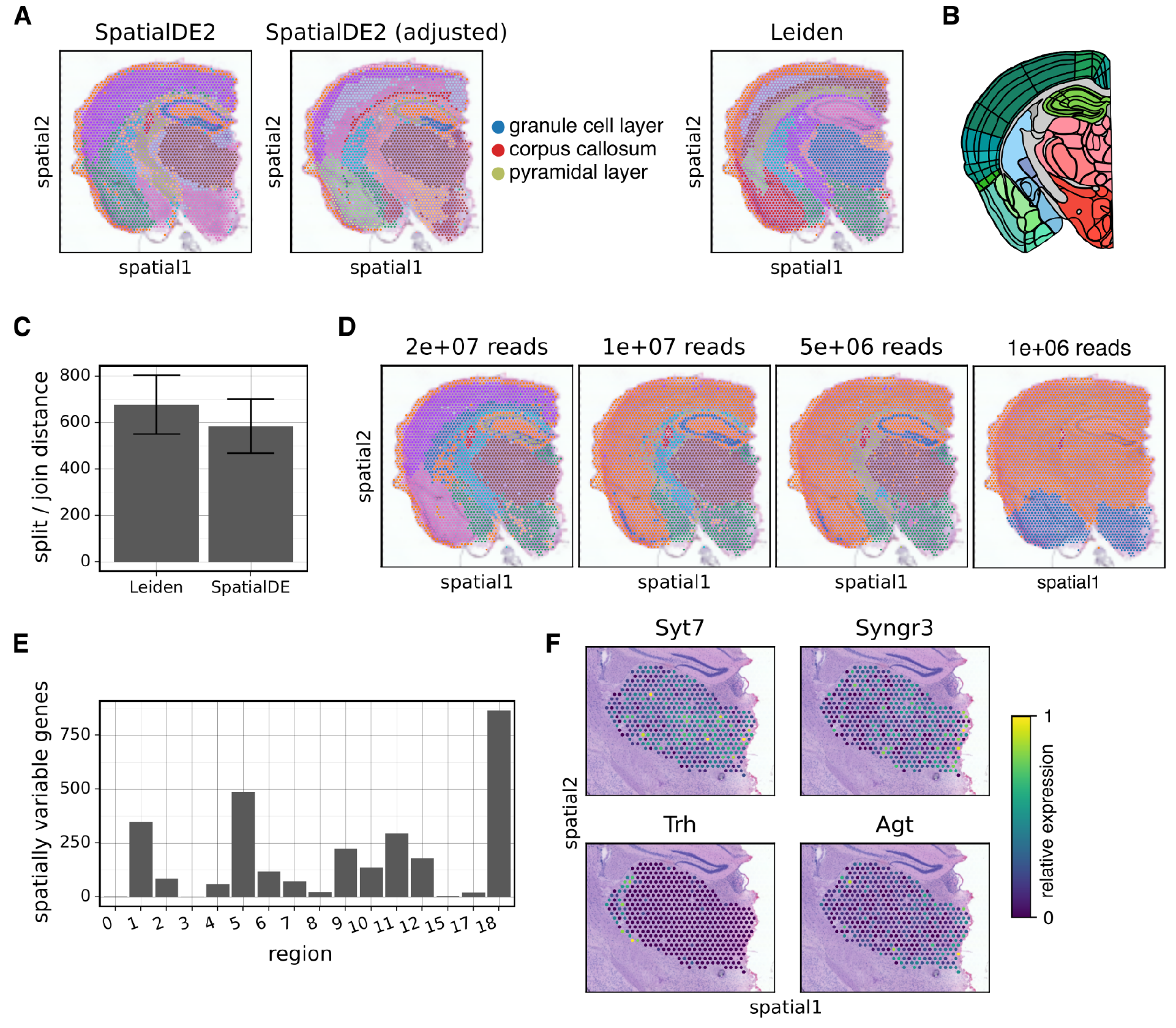
Evaluation of SpatialDE2 tissue region segmentation on the mouse brain. (A) Clustering of mouse brain Visium data with SpatialDE2 using default parameters (left), SpatialDE2 with adjusted parameters (reduced spatial smoothness; middle; c.f. **Supplementary Fig. S2A**) and clustering results obtained using the Leiden workflow (right). Identified tissue regions are annotated in colour. To avoid clutter, only selected regions that are referred to in the main text are labeled in the legend (out of 18 in total) . (B) Corresponding reference annotation of the mouse brain regions obtained from the Allen Brain Atlas (Lein *et al*, 2007). (C) Assessment of the robustness of tissue region segmentations obtained using SpatialDE2 or Leiden clustering. Shown is the concordance across 100 bootstrap experiments, sampling sequencing reads with replacement from the full dataset (N=4.5·10^7^ reads total). Bar height denotes the mean split/join distance compared to the results obtained on the full dataset; error bars denote plus or minus one standard deviation. (D) Assessment of the robustness of the SpatialDE2 tissue region segmentation to downsampling of sequencing reads (out of N=45×10^6 reads total). The total number of sequencing reads in the mouse brain dataset was downsampled to the indicated amount. Shown are the segmentation results. (E) Results from the identification of spatially variable genes within the tissue regions identified from the segmentation step as in (A). Bar heights denote the number of spatially variable genes within each region (FDR<0.1%; out of 12,682 genes assessed). (F) Selected examples of spatially variable genes identified in the thalamus. Shown is normalized relative expression of the indicated genes.

In an attempt to increase spatial resolution, we adjusted the spatial smoothness parameter of SpatialDE2 and the resolution parameter for Leiden clustering. Notably, SpatialDE2 yielded consistent segmentations across a wide range of smoothness parameter settings, providing a meaningful interpolation between coarse- and fine-grained solutions (**Supplementary Fig. S2A**; representative setting shown in **Fig 2A**). As expected, segmentations with a lower smoothness parameter resolved the isocortex layers as well as the corpus callosum (**Fig. 2A**), and further split the hippocampal pyramidal and granule cell layers into separate regions. On the other hand, when varying the resolution parameter of the Leiden workflow, the results were highly variable and lacked intuitive transitions (**Supplementary Fig. S2B**). Furthermore, irrespective of the setting of the resolution parameter, the Leiden algorithm did neither resolve the corpus callosum nor the hippocampal pyramidal/granule cells (**Supplementary Fig. S2B)**.

Analysis of the regions identified by SpatialDE2 in the context of their transcriptomic distance revealed that the solutions obtained by the model respected an appropriate balance between spatial smoothness and transcriptional similarity. In particular, the model assigned the same cluster label to spatially distinct regions if these were transcriptionally very similar. This was particularly apparent in the case of the hippocampal formation and layer 1 of the isocortex, which were assigned to the same cluster by SpatialDE2 (**Supplementary Fig. S2C**).

We also assessed the robustness of SpatialDE2 and Leiden clustering by bootstrapping RNA-seq reads at individual locations (**Fig. 2C**), and we assessed the sensitivity of both methods to variation in sequencing depth (**Fig. 2D, Supplementary Figure S2D**,**E)**. Collectively, these experiments indicated that SpatialDE2 is at least as robust as the Leiden workflow.

Finally, we used the SpatialDE2 spatially variable gene detection module to test for spatially variable genes within the identified tissue regions. This identified up to 867 spatially variable genes (FDR<0.1%; **Fig. 2E**), which were enriched for GO terms related to neurons and nervous tissue, such as “nervous system development” or “synapse” (**Methods, Supplementary Fig. S2D**). Several individual spatially variable genes have known functions in the specific tissue regions, for example Synaptotagmin, Synaptogyrin, Pro-thyrotropin releasing hormone, and Angiotensinogen in the thalamus (**Fig. 2F**). Collectively, these results demonstrate that even within seemingly homogeneous tissue regions, there exist spatial patterns of expression variability at the level of individual genes and pathways, which may help us understand tissue function on a finer spatial scale.

## SpatialDE2 enables fine-grained analyses in complex human tissue

Next, to test SpatialDE2 in a challenging use case, we applied the model to a Visium slide from human endometrium (Garcia-Alonso *et al*, 2021), a highly compartmentalized tissue where cell identity is dependent on spatial context that is just starting to be comprehensively characterized (Garcia-Alonso *et al*, 2021; Wang *et al*, 2020a). Initially, we again applied SpatialDE2 and Leiden clustering to segment tissue regions (**Fig. 3A**). To annotate the regions identified using SpatialDE2, we combined cues from the tissue histology and cell type annotations obtained from computational assignment of reference cell types (using cell2location; (Kleshchevnikov *et al*, 2020)). This identified physiologically meaningful annotations of fine-grained tissue substructures. For example, SpatialDE2 identified regions corresponding to glands and their surrounding areas. In both of these regions, glandular epithelial cells and fibroblasts are present, but in different proportions (**Fig. S3A**). These observed gradual changes in cell type composition are expected and reflect the low resolution of the Visium platform (10 to 50 cells per location). Manual inspection of canonical markers of glandular cells, including *EPCAM* (an epithelial marker) and *PAEP* (a secretory marker typical of glandular cells), confirmed these cell type annotations. Similarly, the muscle cell markers *ACTA2* and *MYLK* were primarily expressed in the myometrium (**Supplementary Fig. S3B**), thus providing additional confidence that SpatialDE2 identified physiologically meaningful tissue regions.

**Figure 3.**
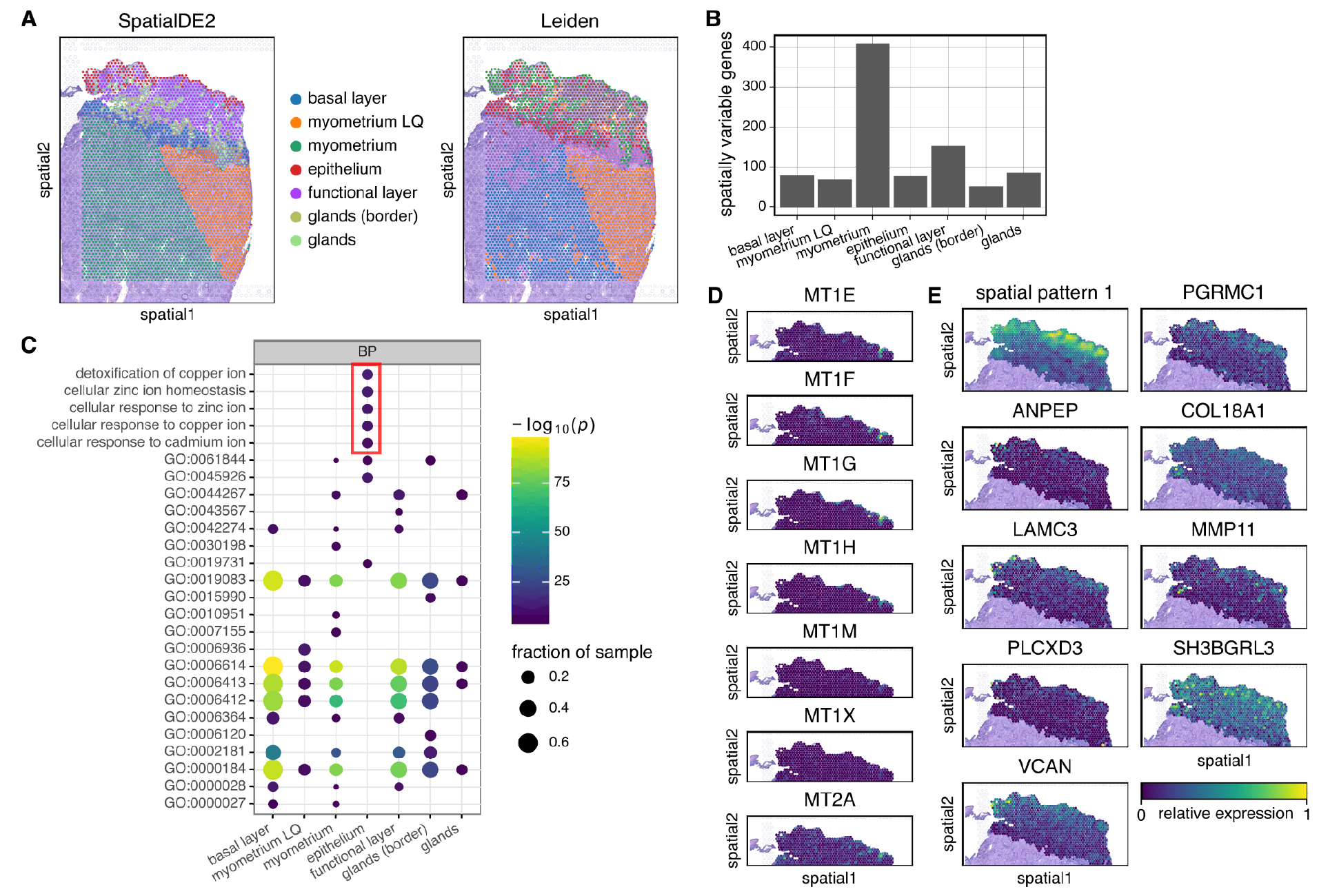
Application of SpatialDE2 to human endometrium tissue. (A) Comparison of tissue segmentation obtained using SpatialDE2 (left) and Leiden clustering (right). SpatialDE2 was used with default settings, and the Leiden resolution parameter was selected to yield a biologically meaningful clustering (see **Supplementary Fig. S3C**) for results using alternative parameters). Myometrium LQ denotes a region of the myometrium with low data quality (Garcia-Alonso *et al*, 2021). (B) Number of spatially variable genes within each region (FDR<0.01%). (C) GO enrichment analysis of spatially variable genes in the named regions of C). GO terms containing metal-binding genes are highlighted. To avoid clutter, only the GO IDs are shown for the other terms. (D) Normalized relative expression of metallothioneins in the functional layer. (E) Normalized relative expression of the progesterone receptor PGRMC1 and other genes belonging to the same spatial pattern (top right)

In contrast to SpatialDE2, the Leiden clustering workflow yielded results that were less concordant to biological expectation. For example, the myometrium regions identified were considerably less smooth than those of SpatialDE2, and functionally distinct uterine regions were assigned to the same cluster: parts of the endometrium basal layer clustered together with a part of the myometrium, another part of the basal layer clustered with parts of the functional layer, and glandular epithelium clustered together with the luminal epithelium (**Fig. 3A**). Critically, while adjustment of the Leiden resolution parameter resulted in partial improvements for some regions - increasing the resolution yielded more fine-grained clustering of the basal and functional endometrial layers and separated the luminal and glandular epithelium – this adjustment simultaneously resulted in overclustering of the myometrium (**Supplementary Fig. S3C**).

Next, we set out to study spatial gene expression within individual tissue regions. The SpatialDE2 test identified between 53 and 409 spatially variable genes per region (FDR<0.001; **Fig. 3B**). GO enrichment analysis identified several GO terms related to metal ions that for spatially variable genes in the epithelium (**Fig. 3C**). Seven of the eight spatially variable genes that were contained in these terms were metallothioneins, which were almost exclusively expressed in a small group of luminal epithelial spots (**Fig. 3D**). The luminal epithelium is enriched in ciliated cells, whose identity depends on the ovarian hormones estrogen and progesterone. Metallothioneins are a feature of the lumenal epithelium in multiple tissues (Danielson *et al*, 1982) have been previously shown to be regulated by progesterone (Burney *et al*, 2007; Slater *et al*, 1988).

We then sought to group genes into patterns of expression using SpatialDE2’s AEH module. This approach is complementary to the segmentation of a tissue into discrete tissue regions, and in particular allows for the identification of smooth transitions. We applied AEH to all spots except those classified as myometrium. We excluded the myometrium for two reasons: It is a relatively homogeneous tissue, and the low-quality region (Garcia-Alonso *et al*, 2021) may interfere with our analysis. This identified five spatial patterns in the endometrium (**Supplementary Figure S3D**). Four of these patterns appeared to recapitulate the distribution of glands in the functional layer, but one pattern (pattern 1) was clearly distinct and contained genes highly expressed in the most luminal part of the functional layer, just below the epithelium (**Fig. 3E**). Among these genes was the progesterone receptor PGRMC1, in agreement with the fact that the functional layer of the endometrium differentiates (decidualizes) in response to hormones.

SpatialDE2 by design takes spatial omics profiles as input, thus rendering the model applicable to a range of different technologies that yield expression estimates that can be explained by a count-based likelihood. This includes both sequencing-based assays but also imaging technologies (see discussion in SpatialDE1 (Svensson *et al*, 2018)). However, in particular for Visium or other technologies that do not achieve true single-cell resolution, the relatively coarse segmentation may result in limitations to assigning specific tissues to clusters, as seen in the endometrium data set with the ‘glands (border)’ cluster. Motivated by this limitation, we set out to explore how SpatialDE2 can be combined with cell type count estimates obtained from computational deconvolution workflows that leverage reference scRNA-seq datasets to estimate cell type abundance. Specifically, we applied SpatialDE2 to the cell type abundance estimates obtained by cell2location on the endometrium dataset (Garcia-Alonso *et al*, 2021). This resulted in a more fine-grained segmentation compared to applying SpatialDE2 directly to gene-counts (**Fig. 4A**). For example, SpatialDE2 now split the basal layer into three distinct regions (numbered 1,5, and 8 in the figure). Spots belonging to these regions were also characterized by distinct clusters in the gene expression space (**Fig. 4B**), indicating that there exist relevant phenotypic differences between these regions. Consistently, we observed marked differences in the cell type composition between these regions; for example region 8 was strongly enriched in epithelial glandular cells, whereas region 1 had more perivascular STEAP4 cells (**Fig 4C**). In general, most regions showed a unique cell type composition (**Supplementary Fig. S4A**), allowing for fine-grained dissection of tissue histology. Finally we note that the Leiden clustering workflow again performed poorly if applied to cell2location output and incorrectly grouped the epithelium and glands as well as parts of the functional and basal layers together, and at the same time severely overclustered the myometrium (**Fig. 4A**).

**Figure 4.**
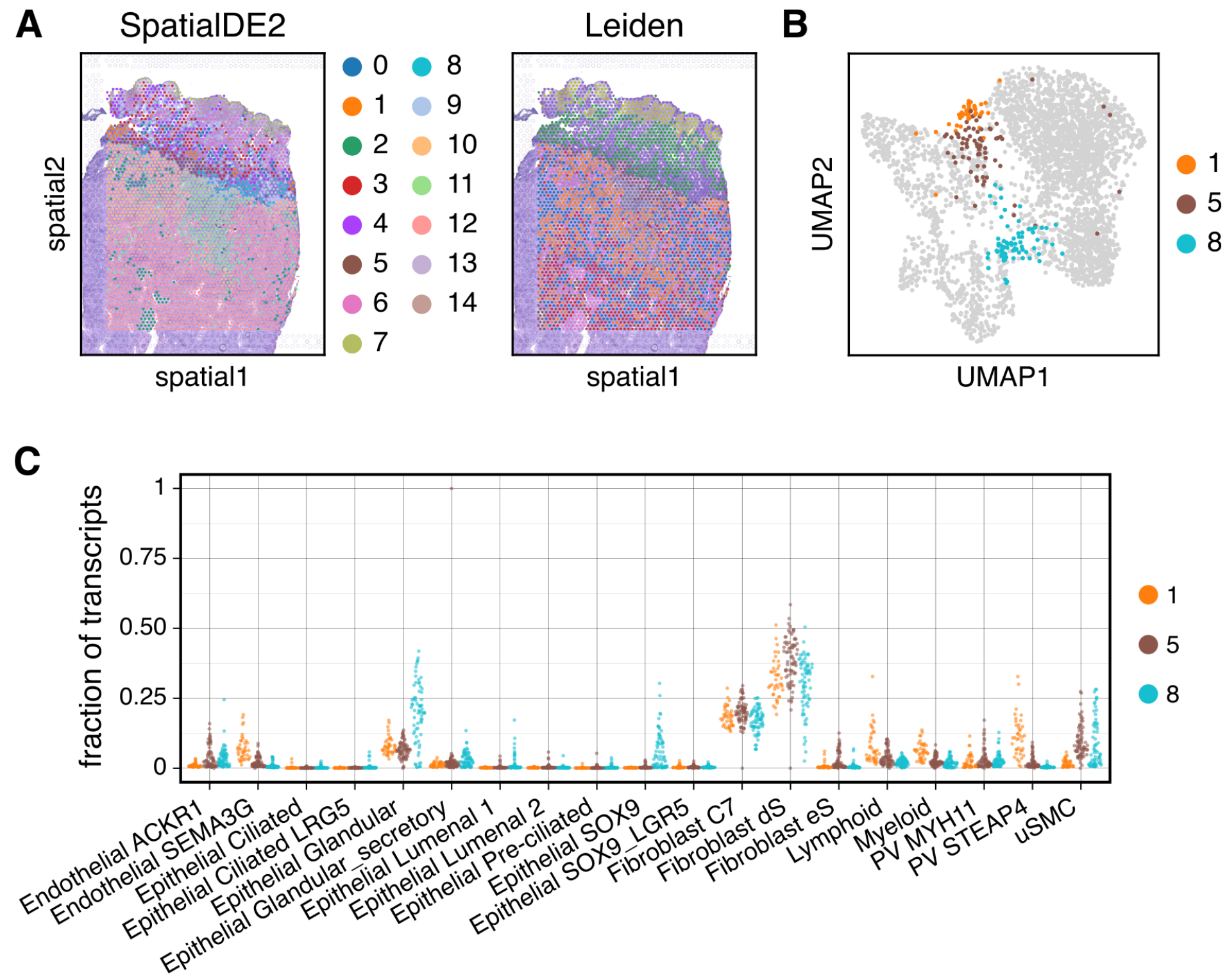
Application of SpatialDE2 to cell type abundance estimates using deconvolution. (A) Comparison of tissue segmentation obtained using SpatialDE2 (left) and Leiden clustering (right) based on cell2location cell type abundance estimates. Number of clusters was determined automatically for SpatialDE2, whereas the Leiden resolution parameter was tuned by hand to yield a biologically meaningful clustering (see **Supplementary Figure S4B** for Leiden clustering with different resolution parameters) (B) UMAP embedding of individual locations based on their expression profiles. Color denotes the cluster labels assigned to one of three basal layer clusters by SpatialDE2. (C) Cell type composition of the three basal layer clusters as in (B). Shown are estimates of cell type fractions from cell2location across individual locations assigned to either cluster 1, 5 or 8 (as in (B)).

## Discussion

Here, we present SpatialDE2, a versatile toolbox for analyzing spatial transcriptomics data. At the core of the method are two analysis modules - region segmentation and spatially variable gene identification. Both modules offer important advantages compared to existing methodology. Moreover, the modules are compatible and can be combined efficiently within one coherent software workflow.

The region segmentation module uniquely allows to account for the spatial structure of tissues, directly operates on RNA count data, and it can be adjusted by a single intuitive parameter. At the same time the module is as efficient as conventional Leiden clustering, which ignores the spatial coordinates. Additionally, SpatialDE2 offers the option of GPU-accelerated computations. Similarly, the spatially variable gene direction module offers orders of magnitude faster computation than previous methods like SpatialDE (Svensson *et al*, 2018) or SPARK (Sun *et al*, 2020), while offering comparable or better statistical power. SpatialDE2 also retains the distinguishing features of these kernel-based methods, such as the ability to test for specific types of expression pattern by designing corresponding kernels.

The two modules work hand in hand, which allows efficient implementation of a workflow consisting of first segmenting the tissue slice followed by detection of spatially variable genes within each region. We illustrated this workflow on the mouse brain, a well-characterized tissue suitable for assessment of performance. On this data set, tissue region segmentation performed comparably to a Leiden clustering workflow and we were able to recover known biology, such as the major brain regions. We also applied the SpatialDE2 workflow to a human endometrium data set, a much less well-characterized tissue. Here, our tissue region segmentation proved superior to Leiden clustering. In addition to recapitulating known biology, SpatialDE2 also generated novel insights into this complex tissue, such as the expression patterns of metallothioneins and several genes spatially co-expressed with the progesterone receptor.

Finally, we showed how SpatialDE2 can be applied downstream of computational deconvolution of spatial transcriptomics based on reference expression profiles, which yielded tissue region segmentation with more fine-grained results, enabling detailed dissection of tissue histology.

Our method is not free of limitations. For one, the size of the kernel and distance matrices used by SpatialDE2 scales with the square of the number of spatial locations. GPUs have only limited memory, which can limit the size of the spatial region that can be analysed using efficient GPU computations. Furthermore, our model is currently designed to be used on one slice at a time. As datasets consisting of multiple sequential tissue sections are becoming more common, future work includes modeling multiple slices at a time.

## Methods

Methods are provided as Supplementary Methods.

## Data Availability

SpatialDE2 is available at https://github.com/PMBio/SpatialDE. Code to reproduce the figures is available at https://github.com/PMBio/spatialde2-paper (note that the data are not included in the repository and need to be downloaded separately from the respective original publications – mouse brain data can be accessed at https://cell2location.cog.sanger.ac.uk/browser.html?shared=tutorial/mouse_brain_visium_data/rawdata/ST8059048/, endometrium data can be accessed at https://www.reproductivecellatlas.org/).

## Acknowledgements

We thank Anna Arutyunyan for kindly sharing the endometrium Visium data and cell2location results and for critically reading the manuscript. We also thank members of the Stegle lab for helpful comments and discussions. The Stegle research group was further supported by core funding from EMBL, the German Cancer Research Center, and the European Commission (ERC project DECODE, 810296).

## Author Contributions

Conceptualization: OS and IK. Formal analysis: IK. Funding acquisition: OS. Methodology: IK and OS. Software: IK. Supervision: OS. Visualization: IK. Writing - original draft: IK and OS. Writing - review & editing: OS, IK, and RVT.

