## Supplementary methods for "SpatialDE2: Fast and localized variance component analysis of spatial transcriptomics"

SpatialDE2 is a flexible framework for modeling spatial transcriptomics data. It comprises two major modules: A tissue region segmentation module and a module for detecting spatially variable genes. In addition, SpatialDE2 contains an automated expression histology (AEH) module, which models expression of a particular gene as drawn from one of a defined number of smooth spatial patterns.
We describe tissue region segmentation in Section 1, spatially variable gene detection in Section 2, and AEH in Section 3. Finally, Section 4 describes applying SpatialDE2 to simulated data and Section 5 deals with analysis of real-world datasets.

#### 1 Tissue region segmentation

The tissue region segmentation module aims to assign a cluster label to each location based on its gene expression profile and the identity of its neighboring locations, with the underlying assumption that neighboring locations likely have the same label, i.e. the segmentation should be spatially smooth.
We implemented tissue segmentation using the infinite hidden markov random field (HMRF) approach[1] adapted to spatial transcriptomics data. In particular, we assume each location belongs to one of $C$ classes. The probability of a location $n$ having label $c$ depends on labeling of location in the neighborhood $\mathcal{N}_n$ of location $n$, and the joint probability follows a Gibbs distribution

$$p(\mathbf{X}) \propto \prod_{n=1}^{N} \prod_{k \in \mathcal{N}_n} \exp\bigl(-E(x_n, x_k)\bigr) \tag{1}$$

where $E(x, y)$ is the pairwise energy of locations $x$ and $y$. Let $x_n$ denote the cluster label of location $n$ and let $\mathbf{x}_n$ and $\mathbf{x}_m$ represent the spatial coordinates of locations $n$ and $m$, respectively. We define

$$E(x_n, x_m) = \begin{cases} \frac{2\sigma^2}{\|\mathbf{x}_n - \mathbf{x}_m\|^2} & x_n \neq x_m \\ 0 & x_n = x_m \end{cases} \tag{2}$$

where $\sigma^2$ is a user-defined hyperparameter. This pairwise energy function promotes spatial smoothness by penalizing different labels for locations in close proximity. Following Chatzis & Tsechpenakis [1] we assume that $p(\mathbf{X})$ can be fully specified as

$$p(\mathbf{X}) = \prod_{n=1}^{N} p(x_n \mid \widehat{\mathbf{x}}_{\mathcal{N}_n}) \tag{3}$$

where

$$p(x_n = c \mid \widehat{\mathbf{x}}_{\mathcal{N}_n}) = \frac{\exp\Bigl(-\sum_{k \in \mathcal{N}_n} E(x_n = c, \widehat{x}_k)\Bigr)}{\sum_{h=1}^{C} \exp\Bigl(-\sum_{k \in \mathcal{N}_n} E(x_n = h, \widehat{x}_k)\Bigr)} \tag{4}$$

and $\widehat{x}_k$ is an estimate of $x_k$.
Gene expression counts $y_{gn}$ for gene $g$ and location $n$ follow a Poisson distribution, with a separate mean $\lambda_{gc}$ for each gene $g$ and class $c$, and differences in sequencing depths between locations are handled by a location-specific scaling factor $S_n$. To enable the model to choose the number of clusters automatically, we use a Dirichlet process prior[1, 2] on the cluster labels. The complete specification of the model is thus

$$\alpha \sim \mathcal{G}\left(\eta_1, \eta_2\right) \tag{5}$$

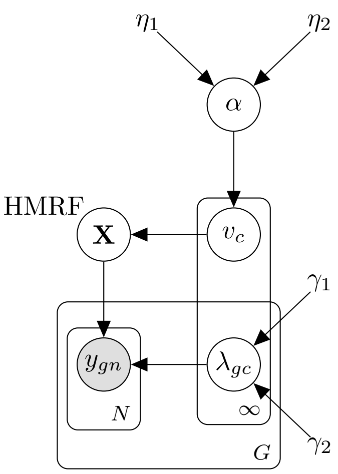


**Figure 1:** Visualization of the tissue region segmentation model in plate notation.

$$v_c \mid \alpha \sim \text{Beta}(1, \alpha) \tag{6}$$
$$x_n \mid \mathbf{v} \sim \text{Categorical}\left(\boldsymbol{\pi}(\mathbf{v})\right) \tag{7}$$
$$\lambda_{gc} \sim \mathcal{G}(\gamma_1, \gamma_2) \tag{8}$$
$$y_{gn} \mid x_n = c, \boldsymbol{\lambda}_g \sim \text{Pois}\left(S_n \lambda_{gc}\right) \tag{9}$$

with

$$\pi_c(\mathbf{v}) = v_c \prod_{c'=1}^{c-1} (1 - v_{c'}) \tag{10}$$

where $\eta_1$, $\eta_2$, $\gamma_1$, and $\gamma_2$ are user-defined hyperparameters. A visualization of the model in plate notation is shown in Fig. 1.
We use variational inference[3, 4] to estimate the posterior distributions. Let $H$ denote the set of hidden variables $\{\mathbf{v}, \alpha, \mathbf{X}, \boldsymbol{\lambda}\}$. We assume that the variational distribution factorizes as $q(H) = q(\mathbf{X})q(\mathbf{v})q(\alpha)q(\boldsymbol{\lambda})$. In the following, let $C$ denote the number of classes. For the generative distribution $C = \infty$, for the variational distribution we follow Blei & Jordan [2] and set $C$ to a fixed integer. Let also $\mathbf{Y} = (y_{gn})_{\substack{g \in [\![1,G]\!] \\ n \in [\![1,N]\!]}}$ denote the complete gene expression count matrix for $G$ genes and $N$ locations.
The complete-data probability is given by

$$p(\mathbf{Y}, \mathbf{X}, \boldsymbol{\lambda}, \mathbf{v}, \alpha) = p(\mathbf{Y}, \mathbf{X} \mid \boldsymbol{\lambda}, \mathbf{v}) p(\mathbf{v} \mid \alpha) p(\boldsymbol{\lambda}) p(\alpha) \tag{11}$$

$p(\mathbf{Y}, \mathbf{X} \mid \boldsymbol{\lambda}, \mathbf{v})$ factorizes:

$$\begin{aligned} p(\mathbf{Y}, \mathbf{X} \mid \boldsymbol{\lambda}, \mathbf{v}) &= \prod_{n=1}^{N} \left( p(x_n \mid x_{\mathcal{N}_n}, \mathbf{v}) \prod_{g=1}^{G} p(y_{gn} \mid x_n, \boldsymbol{\lambda}_\mathbf{g}) \right) \\ &= \prod_{n=1}^{N} \prod_{c=1}^{C} \left( p(x_n = c \mid x_{\mathcal{N}_n})^{\mathbb{1}_{x_n = c}} p(x_n = c \mid \mathbf{v})^{\mathbb{1}_{x_n = c}} \prod_{g=1}^{G} p(y_{gn} \mid \lambda_{gc})^{\mathbb{1}_{x_n = c}} \right) \end{aligned} \tag{12}$$

We now state the variational distributions. $q(\mathbf{X})$ factorizes to

$$\begin{aligned} \log q(x_n = c) \propto \; & \log p(x_n = c \mid x_{\mathcal{N}_n}) + \mathbb{E}_{q(\mathbf{v})} \left[ \sum_{c'=1}^{c-1} \log(1 - v_{c'}) \right] + \mathbb{E}_{q(\mathbf{v})}[\log(v_c)] \\ & + \sum_{g=1}^{G} y_{gn} \, \mathbb{E}_{q(\lambda_{gc})}[\log(\lambda_{gc})] - S_n \sum_{g=1}^{G} \mathbb{E}_{q(\lambda_{gc})}[\lambda_{gc}] \end{aligned} \tag{13}$$

$q(\boldsymbol{\lambda})$ factorizes to

$$q(\lambda_{gc}) = \mathcal{G}\left(\widehat{\gamma}_{1gc}, \widehat{\gamma}_{2gc}\right) \tag{14}$$

where

$$\widehat{\gamma}_{1gc} = \gamma_1 + \sum_{n=1}^{N} q(x_n = c) y_{gn} \tag{15}$$

$$\widehat{\gamma}_{2gc} = \gamma_2 + \sum_{n=1}^{N} q(x_n = c) S_n \tag{16}$$

$$q(\alpha) = \mathcal{G}\left(\eta_1 + C - 1, \widehat{\eta}_2\right) \tag{17}$$

where

$$\widehat{\eta}_2 = \eta_2 - \sum_{c=1}^{C-1} \mathbb{E}_{q(\mathbf{v})}[\log(1 - v_c)] \tag{18}$$

$q(\mathbf{v})$ factorizes to

$$q(v_c) = \mathrm{Beta}\left(\widehat{\alpha}_{1c}, \widehat{\alpha}_{2c}\right) \tag{19}$$

where

$$\widehat{\alpha}_{1c} = 1 + G \sum_{n=1}^{N} q(x_n = c) \tag{20}$$

$$\widehat{\alpha}_{2c} = G \sum_{n=1}^{N} \sum_{i=c+1}^{C} q(x_n = i) + \mathbb{E}_{q(\alpha)}[\alpha] \tag{21}$$

Let us now define

$$\begin{aligned} \widehat{\lambda}_{1gc} &:= \mathbb{E}_{q(\lambda_{gc})}[\lambda_{gc}] \\ &= \frac{\widehat{\gamma}_{1gc}}{\widehat{\gamma}_{2gc}} \end{aligned} \tag{22}$$

$$\begin{aligned} \widehat{\lambda}_{2gc} &:= \mathbb{E}_{q(\lambda_{gc})}[\log \lambda_{gc}] \\ &= \psi(\widehat{\gamma}_{1gc}) - \log \widehat{\gamma}_{2gc} \end{aligned} \tag{23}$$

$$\begin{aligned} \widehat{v}_{1c} &:= \mathbb{E}_{q(v_c)}[v_c] \\ &= \frac{\widehat{\alpha}_{1c}}{\widehat{\alpha}_{1c} + \widehat{\alpha}_{2c}} \end{aligned} \tag{24}$$

$$\begin{aligned} \widehat{v}_{2c} &:= \mathbb{E}_{q(v_c)}[\log v_c] \\ &= \psi(\widehat{\alpha}_{1c}) - \psi(\widehat{\alpha}_{1c} + \widehat{\alpha}_{2c}) \end{aligned} \tag{25}$$

$$\begin{aligned} \widehat{v}_{3c} &:= \mathbb{E}_{q(v_c)}[\log(1 - v_c)] \\ &= \psi(\widehat{\alpha}_{2c}) - \psi(\widehat{\alpha}_{1c} + \widehat{\alpha}_{2c}) \end{aligned} \tag{26}$$

$$\begin{aligned} \widehat{\alpha} &:= \mathbb{E}_{q(\alpha)}[\alpha] \\ &= \frac{\eta_1 + C - 1}{\widehat{\eta}_2} \end{aligned} \tag{27}$$

$$\phi_{nc} := \log p(x_n = c \mid x_{\mathcal{N}_n}) + \sum_{c'=1}^{c-1} \widehat{v}_{3c'} + \widehat{v}_{2c} + \sum_{g=1}^{G} y_{gn} \widehat{\lambda}_{2gc} - S_n \sum_{g=1}^{G} \widehat{\lambda}_{1gc} \tag{28}$$

$$\widehat{\pi}_{nc} := \frac{\exp(\phi_{nc})}{\sum_{c=1}^{C} \exp(\phi_{nc})} \tag{29}$$

where $\psi$ is the digamma function, such that we can now write

$$\widehat{\gamma}_{1gc} = \gamma_1 + \sum_{n=1}^{N} \widehat{\pi}_{nc} y_{gn} \tag{30}$$

$$\widehat{\gamma}_{2gc} = \gamma_2 + \sum_{n=1}^{N} \widehat{\pi}_{nc} S_n \tag{31}$$

$$\widehat{\eta}_2 = \eta_2 - \sum_{c=1}^{C-1} \widehat{v}_{3c} \tag{32}$$

$$\widehat{\alpha}_{1c} = 1 + G\sum_{n=1}^{N}\widehat{\pi}_{nc} \tag{33}$$

$$\widehat{\alpha}_{2c} = G\sum_{n=1}^{N}\sum_{i=c+1}^{C}\widehat{\pi}_{ni} + \widehat{\alpha} \tag{34}$$

Eqs. (22) to (34) define the coupled update equations, with $p(x_n = c \mid x_{\mathcal{N}_n})$ in Eq. (28) given by Eq. (4). Following Chatzis & Tsechpenakis [1], we update the label estimates in each iteration by setting

$$\widehat{x}_n = \underset{c\in[\![1,C]\!]}{\arg\max}\, q(x_n = c) \tag{35}$$

The ELBO is given by

$$\begin{aligned}
\mathcal{L} &= \mathbb{E}_{q(\mathbf{X}),q(\mathbf{v}),q(\boldsymbol{\lambda}),q(\alpha)}[\log p(\mathbf{Y},\mathbf{X},\boldsymbol{\lambda},\mathbf{v},\alpha)] - \mathbb{E}_{q(\mathbf{X}),q(\mathbf{v}),q(\boldsymbol{\lambda}),q(\alpha)}[\log q(\mathbf{X},\boldsymbol{\lambda},\mathbf{v},\alpha)] && (36)\\
&= \mathbb{E}_{q(\mathbf{X}),q(\mathbf{v}),q(\boldsymbol{\lambda})}[\log p(\mathbf{Y},\mathbf{X}\mid\boldsymbol{\lambda},\mathbf{v}) + \mathbb{E}_{q(\mathbf{v}),q(\alpha)}[\log p(\mathbf{v}\mid\alpha)] + \mathbb{E}_{q(\boldsymbol{\lambda})}[\log p(\boldsymbol{\lambda})] + \mathbb{E}_{q(\alpha)}[\log p(\alpha)] \\
&\quad - \mathbb{E}_{q(\mathbf{X})}[\log q(\mathbf{X})] - \mathbb{E}_{q(\boldsymbol{\lambda})}[\log q(\boldsymbol{\lambda})] - \mathbb{E}_{q(\mathbf{v})}[\log q(\mathbf{v})] - \mathbb{E}_{q(\alpha)}[\log q(\alpha)] && (37)\\
&\propto \sum_{n=1}^{N}\sum_{c=1}^{C}\widehat{\pi}_{nc}\log p(x_n = c\mid x_{\mathcal{N}_n}) + \sum_{n=1}^{N}\sum_{c=1}^{C}\widehat{\pi}_{nc}\left(\widehat{v}_{2c} + \sum_{c'=1}^{c-1}\widehat{v}_{3c'}\right) + \sum_{n=1}^{N}\sum_{c=1}^{C}\widehat{\pi}_{nc}\sum_{g=1}^{G}y_{gn}\widehat{\lambda}_{2gc} \\
&\quad - \sum_{n=1}^{N}\sum_{c=1}^{C}\widehat{\pi}_{nc}S_n\sum_{g=1}^{G}\widehat{\lambda}_{1gc} + \sum_{c=1}^{C-1}(\widehat{\alpha}-\widehat{\alpha}_{2c})\widehat{v}_{3c} + \sum_{g=1}^{G}\sum_{c=1}^{C}(\gamma_1-\widehat{\gamma}_{1gc})\widehat{\lambda}_{2gc} + \sum_{g=1}^{G}\sum_{c=1}^{C}(\widehat{\gamma}_{2gc}-\gamma_2)\widehat{\lambda}_{1gc} \\
&\quad - \sum_{n=1}^{N}\sum_{c=1}^{C}\widehat{\pi}_{nc}\phi_{nc} + \sum_{n=1}^{N}\log\left(\sum_{c=1}^{C}\exp(\phi_{nc})\right) - \sum_{g=1}^{G}\sum_{c=1}^{C}\left(\widehat{\gamma}_{1gc}\log\widehat{\gamma}_{2gc} - \log\Gamma\left(\widehat{\gamma}_{1gc}\right)\right) && (38)\\
&\quad - \sum_{c=1}^{C-1}(\widehat{\alpha}_{1c}-1)\widehat{v}_{2c} + \sum_{c=1}^{C-1}\log B(\widehat{\alpha}_{1c},\widehat{\alpha}_{2c}) - (\eta_1 + C)\log\widehat{\eta}_2 + (\widehat{\eta}_2-\eta_2)\widehat{\alpha}
\end{aligned}$$

### 2 Spatially variable gene detection

Suppose that there are multiple sources of expression variability for a particular gene. Some of these sources may depend on spatial location and induce correlations between locations in close spatial proximity, while others may depend on some other covariates or represent random noise. Detecting spatially variable genes entails finding genes with a statistically significant non-zero amount of structured spatial variability.

#### 2.1 General model and omnibus test

We use the Gaussian process framework to model structured spatial variability. In particular, we propose the following general model for spatial transcriptomics data:

$$\begin{aligned}
\mathbf{e} &\sim \mathcal{N}\left(\mu\mathbf{1}, \sigma_1\boldsymbol{\Sigma}_1 + \cdots + \sigma_n\boldsymbol{\Sigma}_n\right) \\
\mathbf{y}\mid\mathbf{e} &\sim \text{Pois}\left(\mathbf{s}\odot\exp(\mathbf{e})\right)
\end{aligned} \tag{39}$$

where $\mathbf{y} = \mathbf{Y}_{:,g}$ is the measured expression of a gene, $\mathbf{s}$ is a vector of location/cell-specific scaling factors and $\odot$ denotes the Hadamard product. To test the null hypothesis $H_0 : \sigma_1 = \cdots = \sigma_k = 0$, $k \le n$, we employ a variance component score test based on Zhang & Lin [5], which is computationally efficient, since it only requires the model under the null hypothesis to be estimated. The score test of Zhang & Lin [5] tests the null hypothesis $H_0 : \sigma_i = 0$ for some $i$, but their derivation is easily extended to yield an omnibus test capable of testing our generalized null hypothesis.

Let us phrase the problem as a generalized linear model (GLMM) and for the sake of notational simplicity assume without loss of generality that $k = n$. Let $\mathbf{a}_1 \sim \mathcal{N}\left(\mathbf{0},\sigma_1\boldsymbol{\Sigma}_1\right),\ldots,\mathbf{a}_k \sim \mathcal{N}\left(\mathbf{0},\sigma_k\boldsymbol{\Sigma}_k\right)$. We can define $\mathbf{a} \coloneqq \sum_{i=1}^{k}\mathbf{a}_i$ and $\boldsymbol{\sigma} = (\sigma_1,\ldots,\sigma_k)^{\mathrm{T}}$, such that $\mathbf{a} \sim \mathcal{N}\left(\mathbf{0},\sum_{i=1}^{k}\sigma_i\boldsymbol{\Sigma}_i\right)$. The observations $y_i$ are assumed to have means $\mathbb{E}[y_i] = \mu_i$ and variances $\mathbb{V}\text{ar}[y_i] = \phi\omega_i^{-1}v(\mu_i)$, where $\phi$ is a dispersion parameter, $\omega_i$ is a known prior weight, and $v(\cdot)$ is a variance function. Let $\mathbf{X}$ be a fixed effect design matrix and consider the GLMM

$$g(\boldsymbol{\mu}) = \mathbf{X}\boldsymbol{\beta} + \mathbf{a} \tag{40}$$

where $\boldsymbol{\beta}$ is a fixed effect and $g(\cdot)$ is a link function. Let further $\boldsymbol{\Delta} := \operatorname{diag}(g'(\boldsymbol{\mu}))$ and $\mathbf{W} := \operatorname{diag}(\mathbf{w})$ with $w_i := \phi\omega_i^{-1}v(\mu_i)(g'(\mu_i))^2$. Then the score vector for the null hypothesis $H_0 : \sigma_1 = \cdots = \sigma_k = 0$ has components

$$\mathcal{U}_{\sigma_i} \approx \frac{1}{2}\left((\mathbf{y}-\boldsymbol{\mu}^{(0)})^{\mathrm{T}}\boldsymbol{\Delta}\mathbf{W}^{-1}\boldsymbol{\Sigma}_i\mathbf{W}^{-1}\boldsymbol{\Delta}(\mathbf{y}-\boldsymbol{\mu}^{(0)}) - \operatorname{tr}(\mathbf{P}\boldsymbol{\Sigma}_i)\right)\Bigg|_{\boldsymbol{\beta}=\widehat{\boldsymbol{\beta}}} \tag{41}$$

where $\boldsymbol{\mu}^{(0)}$ satisfies the null parametric generalized mixed model $g(\boldsymbol{\mu}^{(0)}) = \mathbf{X}\boldsymbol{\beta}$, $\widehat{\boldsymbol{\beta}}$ is a BLUP estimate of $\boldsymbol{\beta}$ and we have defined $\mathbf{P} := \mathbf{W}^{-1} - \mathbf{W}^{-1}\mathbf{X}(\mathbf{X}^{\mathrm{T}}\mathbf{W}^{-1}\mathbf{X})^{-1}\mathbf{X}^{\mathrm{T}}\mathbf{W}^{-1}$. Eq. (41) is identical to Eq. (16) of Zhang & Lin [5].

Writing $\mathcal{U}_{\sigma_i} = U_{\sigma_i} - e_{\sigma_i}$, where $U_{\sigma_i}$ and $e_{\sigma_i}$ are the first and second terms in Eq. (41), we define the test statistic

$$\begin{aligned} U_{\boldsymbol{\sigma}} &= \sum_{i=1}^{k} U_{\sigma_i} \\ &= \frac{1}{2}(\mathbf{y}-\boldsymbol{\mu}^{(0)})^{\mathrm{T}}\boldsymbol{\Delta}\mathbf{W}^{-1}\left(\sum_{i=1}^{k}\boldsymbol{\Sigma}_i\right)\mathbf{W}^{-1}\boldsymbol{\Delta}(\mathbf{y}-\boldsymbol{\mu}^{(0)}) \end{aligned} \tag{42}$$

$U_{\boldsymbol{\sigma}}$ asymptotically follows a mixture of $\chi^2$ distributions that we approximate using the Satterthwaite method as in Zhang & Lin [5] to calculate a p-value.

#### 2.2 Prior work

The above model is a generalization of previous methods like SpatialDE[6], SVCA[7], and SPARK[8], containing the latter as special cases. The SVCA model contains four variance components, representing intrinsic, environmental, and cell-cell interactions as well as random noise, respectively. SpatialDE and SPARK are even simpler in that they use two variance components representing structured spatial variability and random noise, respectively.

#### 2.3 SpatialDE2 implementation: Optimizing computational efficiency

The special case implemented by SpatialDE and SPARK permits a further simplification: Under the null hypothesis, only the noise variance component remains and the observations essentially follow an overdispersed Poisson distribution. To avoid fitting a GLMM in this simple case, we implemented the SpatialDE null model as a Negative Binomial with constant mean and an overdispersion parameter.

Spatial variability is modeled by a kernel covariance matrix $\mathbf{K}$ with $K_{i,j} = k(\mathbf{x}_i, \mathbf{x}_j)$, where $k$ is a kernel function and $\mathbf{x}_i$ and $\mathbf{x}_j$ are spatial coordinates of the $i$-th and $j$-th data points, respectively. The model then becomes

$$\begin{aligned} \mathbf{e} &\sim \mathcal{N}(\mu\mathbf{1}, \sigma_s\mathbf{K} + \sigma_n\mathbf{I}) \\ \mathbf{y} \mid \mathbf{e} &\sim \operatorname{Pois}(\mathbf{s} \odot \exp(\mathbf{e})) \end{aligned} \tag{43}$$

Kernel functions include the squared exponental kernel function $k_{\mathrm{sqexp}}(\mathbf{x},\mathbf{y}) = \exp\left(-\frac{\|\mathbf{x}-\mathbf{y}\|_2^2}{2l^2}\right)$, the periodic (cosine) kernel function $k_{\cos}(\mathbf{x},\mathbf{y}) = \cos\left(\frac{2\pi\sum_k(x_k-y_k)}{l}\right)$, and the linear kernel function $k_{\mathrm{lin}}(\mathbf{x},\mathbf{y}) = \mathbf{x}^{\mathrm{T}}\mathbf{y}$. While not required for fitting the null model, the covariance (kernel) matrix of the spatial variance component is however still required for the significance test. This poses a problem, since some kernel functions include hyperparameters that are not known *a priori*. For the squared exponential and the cosine kernels, the unknown hyperparameter is the lengthscale $l$, indicating the typical distance (squared exponential) or period length (cosine) of the spatial variation. Moreover, the particular kernel function itself is not known *a priori*. SpatialDE approached these issues by first fitting the alternative model for each kernel function and hyperparameter separately, selecting the best fiting model, and performing the significance test for this model only. This approach negates the computational advantage of the score test, which only requires the simpler null model to be fitted. We therefore follow SPARK's approach of performing the score test for several pre-specified kernels with different structures and lengthscales, but implement an additional optimization: Whereas SPARK tests each kernel individually and combines the p-values from all tests to select spatially variable genes, we reinterpret this problem as an omnibus test. Let $\mathbf{K}_1, \ldots, \mathbf{K}_k$ denote the kernels to be tested, then we can write the model as

$$\begin{aligned} \mathbf{e} &\sim \mathcal{N}(\mu\mathbf{1}, \sigma_1\mathbf{K}_1 + \cdots + \sigma_k\mathbf{K}_k + \sigma_n\mathbf{I}) \\ \mathbf{y} \mid \mathbf{e} &\sim \operatorname{Pois}(\mathbf{s} \odot \exp(\mathbf{e})) \end{aligned} \tag{44}$$

with $H_0 : \sigma_1 = \cdots = \sigma_k = 0$ and apply the omnibus test described above. Computationally, this is equivalent to performing a single score test on the sum of all kernel matrices. A single score test requires several expensive matrix operations to be performed, scaling as $\mathcal{O}(N^2)$ when optimally implemented, where $N$ is the number of observations. SPARK's approach in addition scales linearly with the number of kernels $K$ with a total complexity of $\mathcal{O}(KN^2)$, while our implementation's complexity is independent of the number of kernels. Using ten different kernels by default, our implementation is therefore ten times faster than SPARK. We implemented both model fitting and significance testing in Tensorflow[9–11], taking advantage of GPU acceleration and further increasing speed by orders of magnitude.

**Table S1:** Features of several spatial transcriptomics analysis packages based on Gaussian processes

| | SpatialDE2 | SpatialDE[6] | SVCA[7] | SPARK[8] |
|---|---|---|---|---|
| spatially variable gene detection | yes | yes | no | yes |
| spatial variance decomposition | yes | no | yes | no |
| likelihood model | count-based | Gaussian approximation | Gaussian approximation | count-based |
| statistical test | score test | likelihood ratio test w/ theoretical $\chi^2$ distribution | likelihood ratio test w/ emprirical $\chi^2$ distribution | score test |
| testing multiple covariances | test separately and combine p-values or omnibus test | fit all, test only best fit | only squared exponential supported | test separately and combine p-values |
| GPU acceleration | yes | no | no | no |
| Language | Python | Python | Python | R |

### 3 Automated Expression Histology (AEH)

Automated expression histology aims to group genes into a small number of groups, such that all genes in a group share a common spatial expression pattern. This is achieved by fitting a Gaussian process mixture model, with several Gaussian processes representing smooth spatial patterns, and expression of each gene drawn from one Gaussian process with added noise. AEH was first introduced by SpatialDE[6].

We extend SpatialDE's AEH implementation by a Dirichlet process prior[1, 2] to automatically select the number of spatial patterns. In detail, we assume that there are $C$ spatial expression patterns $\boldsymbol{\mu}_c$ in the data set, and each gene belongs to one of these patterns. We assume that expression $\mathbf{y}_g$ of gene $g$ has been normalized to be approximately Gaussian, and we put a Gaussian process prior on $\boldsymbol{\mu}_c$. All genes are assumed to share a common noise variance $\sigma$. The complete specification of the model is thus

$$
\begin{aligned}
\alpha &\sim \mathcal{G}\left(\eta_1, \eta_2\right) && (45)\\
v_c \mid \alpha &\sim \operatorname{Beta}\left(1, \alpha\right) && (46)\\
x_g \mid \mathbf{v} &\sim \operatorname{Categorical}\left(\boldsymbol{\pi}(\mathbf{v})\right) && (47)\\
\sigma &\sim \operatorname{Inv}\mathcal{G}\left(\gamma_1, \gamma_2\right) && (48)\\
\boldsymbol{\mu}_c &\sim \mathcal{N}\left(\mathbf{0}, \boldsymbol{\Sigma}_c\right) && (49)\\
\mathbf{y}_g \mid x_g = c, \boldsymbol{\mu}, \boldsymbol{\sigma} &\sim \mathcal{N}\left(\boldsymbol{\mu}_c, \sigma\mathbf{I}\right) && (50)
\end{aligned}
$$

with

$$\pi_c(\mathbf{v}) = v_c \prod_{c'=1}^{c-1} (1 - v_{c'}) \tag{51}$$

where $x_g$ is the pattern label of gene $g$. A visualization of the model in plate notation is shown in Fig. 2.
The complete-data probability is given by

$$p(\mathbf{Y}, \mathbf{x}, \boldsymbol{\mu}, \sigma, \mathbf{v}, \alpha) = p(\mathbf{Y} \mid \boldsymbol{\mu}, \sigma, \mathbf{x}) p(\boldsymbol{\mu}) p(\sigma) p(\mathbf{x} \mid \mathbf{v}) p(\mathbf{v} \mid \alpha) p(\alpha) \tag{52}$$

with

$$p(\mathbf{Y} \mid \boldsymbol{\mu}, \sigma, \mathbf{x}) = \prod_{g=1}^{G} \prod_{c=1}^{\infty} \mathcal{N}\left(\mathbf{y}_g | \boldsymbol{\mu}_c, \sigma\mathbf{I}\right)^{\mathbb{1}_{x_g = c}} \tag{53}$$

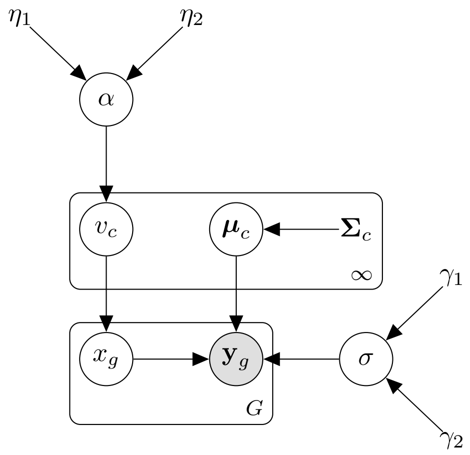


**Figure 2:** Visualization of the AEH model in plate notation.

$$p(\sigma) = \prod_{c=1}^{\infty} \text{Inv}\mathcal{G}\left(\sigma|\gamma_1, \gamma_2\right) \tag{54}$$

$$p(\boldsymbol{\mu}) = \prod_{c=1}^{\infty} \mathcal{N}\left(\boldsymbol{\mu}_c|\mathbf{0}, \boldsymbol{\Sigma}_c\right) \tag{55}$$

$$p(\mathbf{x} \mid \mathbf{v}) = \prod_{g=1}^{G} \prod_{c=1}^{\infty} \pi_c^{\mathbb{1}_{x_g=c}} \tag{56}$$

$$p(\mathbf{v} \mid \alpha) = \prod_{c=1}^{\infty} \text{Beta}\left(v_c|1, \alpha\right) \tag{57}$$

We use variational inference[3, 4] to estimate the posterior distributions. Let $H$ denote the set of hidden variables $\{\mathbf{v}, \mathbf{x}, \alpha, \boldsymbol{\mu}, \boldsymbol{\sigma}\}$. We assume that the variational distribution factorizes as $q(H) = q(\mathbf{v})q(\mathbf{x})q(\alpha)q(\boldsymbol{\mu})q(\sigma)$. In the following, let $C$ denote the number of classes. For the generative distribution $C = \infty$, for the variational distribution we follow Blei & Jordan [2] and set $C$ to a fixed integer.
The variational distributions are given by

$$\begin{aligned} \log q(x_g = c) \propto\ & \mathbb{E}_{q(\mathbf{v})}\left[\sum_{c'=1}^{c-1} \log(1 - v_{c'})\right] + \mathbb{E}_{q(\mathbf{v})}[\log(v_c)] - \frac{N}{2}\,\mathbb{E}_{q(\sigma)}\left[\log \sigma\right] - \frac{1}{2}\,\mathbb{E}_{q(\sigma)}\left[\frac{1}{\sigma}\right] \mathbf{y}_g^{\mathrm{T}} \mathbf{y}_g \\ & + \mathbb{E}_{q(\sigma)}\left[\frac{1}{\sigma}\right] \mathbf{y}_g^{\mathrm{T}}\, \mathbb{E}_{q(\boldsymbol{\mu})}[\boldsymbol{\mu}_c] - \frac{1}{2}\,\mathbb{E}_{q(\boldsymbol{\sigma})}\left[\frac{1}{\sigma}\right] \mathbb{E}_{q(\boldsymbol{\mu})}[\boldsymbol{\mu}_c^{\mathrm{T}} \boldsymbol{\mu}_c] \end{aligned} \tag{58}$$

$$q(\alpha) = \mathcal{G}\left(\eta_1 + C - 1, \widehat{\eta}_2\right) \tag{59}$$

with

$$\widehat{\eta}_2 = \eta_2 - \sum_{c=1}^{C-1} \mathbb{E}_{q(\mathbf{v})}[\log(1 - v_c)] \tag{60}$$

$q(\mathbf{v})$ factorizes to

$$q(v_c) = \text{Beta}\left(\widehat{\alpha}_{1c}, \widehat{\alpha}_{2c}\right) \tag{61}$$

with

$$\widehat{\alpha}_{1c} = 1 + \sum_{g=1}^{G} q(x_g = c) \tag{62}$$

$$\widehat{\alpha}_{2c} = \sum_{g=1}^{G} \sum_{i=c+1}^{C} q(x_g = i) + \mathbb{E}_{q(\alpha)}[\alpha] \tag{63}$$

$q(\boldsymbol{\mu})$ factorizes to

$$q(\boldsymbol{\mu}_c) = \mathcal{N}\left(\widehat{\boldsymbol{\mu}}_c, \widehat{\boldsymbol{\Sigma}}_c\right) \tag{64}$$

with

$$\widehat{\boldsymbol{\Sigma}}_c = \left( \mathbb{E}_{q(\sigma)} \left[ \frac{1}{\sigma} \right] \sum_{g=1}^{G} q(x_g = c) \mathbf{I} + \boldsymbol{\Sigma}_c^{-1} \right)^{-1} \tag{65}$$

$$\widehat{\boldsymbol{\mu}}_c = \left( \mathbb{E}_{q(\sigma)} \left[ \frac{1}{\sigma} \right] \sum_{g=1}^{G} q(x_g = c) \mathbf{I} + \boldsymbol{\Sigma}_c^{-1} \right)^{-1} \mathbb{E}_{q(\sigma)} \left[ \frac{1}{\sigma} \right] \sum_{g=1}^{G} q(x_g = c) \mathbf{y}_g \tag{66}$$

$$= \widehat{\boldsymbol{\Sigma}}_c \, \mathbb{E}_{q(\sigma)} \left[ \frac{1}{\sigma} \right] \sum_{g=1}^{G} q(x_g = c) \mathbf{y}_g \tag{67}$$

$$q(\sigma) = \mathrm{Inv}\mathcal{G} \left( \widehat{\gamma}_1, \widehat{\gamma}_2 \right) \tag{68}$$

with

$$\widehat{\gamma}_1 = \gamma_1 + \frac{NG}{2} \tag{69}$$

$$\widehat{\gamma}_2 = \gamma_2 + \frac{1}{2} \left( \sum_{g=1}^{G} \mathbf{y}_g^{\mathrm{T}} \mathbf{y}_g - 2 \sum_{c=1}^{C} \left( \sum_{g=1}^{G} q(x_g = c) \mathbf{y}_g \right)^{\mathrm{T}} \widehat{\boldsymbol{\mu}}_c + \sum_{c=1}^{C} \sum_{g=1}^{G} q(x_g = c) \left( \widehat{\boldsymbol{\mu}}_c^{\mathrm{T}} \widehat{\boldsymbol{\mu}}_c + \mathrm{Tr}\left( \widehat{\boldsymbol{\Sigma}}_c \right) \right) \right) \tag{70}$$

Let us now define

$$\widehat{v}_{1c} \coloneqq \mathbb{E}_{q(v_c)}[v_c] = \frac{\widehat{\alpha}_{1c}}{\widehat{\alpha}_{1c} + \widehat{\alpha}_{2c}} \tag{71}$$

$$\widehat{v}_{2c} \coloneqq \mathbb{E}_{q(v_c)}[\log v_c] = \psi(\widehat{\alpha}_{1c}) - \psi(\widehat{\alpha}_{1c} + \widehat{\alpha}_{2c}) \tag{72}$$

$$\widehat{v}_{3c} \coloneqq \mathbb{E}_{q(v_c)}[\log(1 - v_c)] = \psi(\widehat{\alpha}_{2c}) - \psi(\widehat{\alpha}_{1c} + \widehat{\alpha}_{2c}) \tag{73}$$

$$\widehat{\alpha} \coloneqq \mathbb{E}_{q(\alpha)}[\alpha] = \frac{\eta_1 + C - 1}{\widehat{\eta}_2} \tag{74}$$

$$\widehat{\sigma} \coloneqq \mathbb{E}_{q(\sigma)} \left[ \frac{1}{\sigma} \right] = \frac{\widehat{\gamma}_1}{\widehat{\gamma}_2} \tag{75}$$

$$\widehat{l} \coloneqq \mathbb{E}_{q(\sigma)} \left[ \log \sigma \right] = \log \widehat{\gamma}_2 - \psi(\widehat{\gamma}_1) \tag{76}$$

$$\phi_{gc} \coloneqq \sum_{c'=1}^{c-1} \widehat{v}_{3c'} + \widehat{v}_{2c} - \frac{N}{2} \widehat{l} - \frac{1}{2} \widehat{\sigma} \left( \mathbf{y}_g^{\mathrm{T}} \mathbf{y}_g - 2 \mathbf{y}_g^{\mathrm{T}} \widehat{\boldsymbol{\mu}}_c + \widehat{\boldsymbol{\mu}}_c^{\mathrm{T}} \widehat{\boldsymbol{\mu}}_c + \mathrm{Tr}\left( \widehat{\boldsymbol{\Sigma}}_c \right) \right) \tag{77}$$

$$\widehat{\pi}_{gc} \coloneqq \frac{\exp\left(\phi_{gc}\right)}{\sum_{c=1}^{C} \exp\left(\phi_{gc}\right)} \tag{78}$$

$$N_c \coloneqq \sum_{g=1}^{G} \widehat{\pi}_{gc} \tag{79}$$

$$\overline{\mathbf{y}}_c \coloneqq \sum_{g=1}^{G} \widehat{\pi}_{gc} \mathbf{y}_g \tag{80}$$

where $\psi$ is the digamma function, such that we can write

$$\widehat{\alpha}_{1c} = 1 + N_c \tag{81}$$

$$\widehat{\alpha}_{2c} = \sum_{g=1}^{G} \sum_{i=c+1}^{C} \widehat{\pi}_{gi} + \widehat{\alpha} \tag{82}$$

$$\widehat{\eta}_2 = \eta_2 - \sum_{c=1}^{C-1} \widehat{v}_{3c} \tag{83}$$

$$\begin{aligned}\widehat{\boldsymbol{\Sigma}}_c &= \left(\widehat{\sigma} N_c \mathbf{I} + \boldsymbol{\Sigma}_c^{-1}\right)^{-1} \\ &= \boldsymbol{\Sigma}_c \left(\widehat{\sigma} N_c \boldsymbol{\Sigma}_c + \mathbf{I}\right)^{-1} \quad \text{using eq. 161 from the Matrix Cookbook} \\ &= \left(\widehat{\sigma} N_c \boldsymbol{\Sigma}_c + \mathbf{I}\right)^{-\mathrm{T}} \boldsymbol{\Sigma}_c^{\mathrm{T}} \quad \text{using symmetry of } \widehat{\boldsymbol{\Sigma}}_c \\ &= \left(\widehat{\sigma} N_c \boldsymbol{\Sigma}_c + \mathbf{I}\right)^{-1} \boldsymbol{\Sigma}_c \quad \text{using symmetry of } \boldsymbol{\Sigma}_c\end{aligned} \tag{84}$$

$$\widehat{\boldsymbol{\mu}}_c = \widehat{\boldsymbol{\Sigma}}_c \widehat{\sigma} \overline{\mathbf{y}}_c \tag{85}$$

$$\widehat{\gamma}_2 = \gamma_2 + \frac{1}{2}\left(\sum_{g=1}^{G} \mathbf{y}_g^{\mathrm{T}} \mathbf{y}_g - 2\sum_{c=1}^{C} \overline{\mathbf{y}}_c^{\mathrm{T}} \widehat{\boldsymbol{\mu}}_c + \sum_{c=1}^{C} N_c \widehat{\boldsymbol{\mu}}_c^{\mathrm{T}} \widehat{\boldsymbol{\mu}}_c + \sum_{c=1}^{C} N_c \operatorname{Tr}\left(\widehat{\boldsymbol{\Sigma}}_c\right)\right) \tag{86}$$

The ELBO is given by

$$\mathcal{L} = \mathbb{E}_{q(\boldsymbol{\mu}),q(\sigma),q(\mathbf{x}),q(\mathbf{v}),q(\alpha)}[\log p(\mathbf{Y}, \mathbf{x}, \boldsymbol{\mu}, \sigma, \mathbf{v}, \alpha)] - \mathbb{E}_{q(\boldsymbol{\mu}),q(\sigma),q(\mathbf{x}),q(\mathbf{v}),q(\alpha)}[\log q(\mathbf{x}, \boldsymbol{\mu}, \sigma, \mathbf{v}, \alpha)] \tag{87}$$

$$\begin{aligned}&= \mathbb{E}_{q(\boldsymbol{\mu}),q(\sigma),q(\mathbf{x})}[\log p(\mathbf{Y} \mid \boldsymbol{\mu}, \sigma, \mathbf{x})] + \mathbb{E}_{q(\boldsymbol{\mu})}[\log p(\boldsymbol{\mu})] + \mathbb{E}_{q(\mathbf{x},q(\mathbf{v}))}[\log p(\mathbf{x} \mid \mathbf{v})] \\ &\quad + \mathbb{E}_{q(\mathbf{v}),q(\alpha)}[\log p(\mathbf{v} \mid \alpha)] + \mathbb{E}_{q(\alpha)}[\log p(\alpha)] - \mathbb{E}_{q(\mathbf{x})}[\log q(\mathbf{x})] - \mathbb{E}_{q(\mathbf{v})}[\log q(\mathbf{v})] \\ &\quad - \mathbb{E}_{q(\boldsymbol{\mu})}[\log q(\boldsymbol{\mu})] - \mathbb{E}_{q(\sigma)}[\log q(\sigma)] - \mathbb{E}_{q(\alpha)}[\log q(\alpha)]\end{aligned} \tag{88}$$

$$\begin{aligned}&\propto -\frac{NG}{2}\widehat{l} - \frac{1}{2}\widehat{\sigma}\left(\sum_{g=1}^{G} \mathbf{y}_g^{\mathrm{T}} \mathbf{y}_g - 2\sum_{c=1}^{C} \overline{\mathbf{y}}_c^{\mathrm{T}} \widehat{\boldsymbol{\mu}}_c + \sum_{c=1}^{C} N_c \widehat{\boldsymbol{\mu}}_c^{\mathrm{T}} \widehat{\boldsymbol{\mu}}_c + \sum_{c=1}^{C} N_c \operatorname{Tr}\left(\widehat{\boldsymbol{\Sigma}}_c\right)\right) \\ &\quad - \frac{1}{2}\sum_{c=1}^{C}\left(\widehat{\boldsymbol{\mu}}_c^{\mathrm{T}} \widehat{\sigma}(\widehat{\sigma} N_c \boldsymbol{\Sigma}_c + \mathbf{I})^{-1} \overline{\mathbf{y}}_c + \operatorname{Tr}\left((\widehat{\sigma}_c N_c \boldsymbol{\Sigma}_c + \mathbf{I})^{-1}\right)\right) - \frac{1}{2}\sum_{c=1}^{C} \log|\widehat{\sigma}_c N_c \boldsymbol{\Sigma}_c + \mathbf{I}| + \sum_{c=1}^{C-1} N_c \widehat{v}_{2c} \\ &\quad + \sum_{c=1}^{C} N_c \sum_{c'=1}^{c-1} \widehat{v}_{3c'} + \sum_{c=1}^{C-1} (\widehat{\alpha} - \widehat{\alpha}_{2c}) \widehat{v}_{3c} - \sum_{g=1}^{G}\sum_{c=1}^{C} \widehat{\pi}_{gc} \phi_{gc} + \sum_{g=1}^{G} \log\left(\sum_{c=1}^{C} \exp(\phi_{gc})\right) - \widehat{\gamma}_1(1 - \log \widehat{\gamma}_2) \\ &\quad + \log\Gamma(\widehat{\gamma}_1) + (\widehat{\gamma}_1 + 1)\widehat{l} - \sum_{c=1}^{C-1} (\widehat{\alpha}_{1c} - 1)\widehat{v}_{2c} + \sum_{c=1}^{C-1} \log B(\widehat{\alpha}_{1c}, \widehat{\alpha}_{2c}) - (\eta_1 + C)\log \widehat{\eta}_2 + (\widehat{\eta}_2 - \eta_2)\widehat{\alpha}\end{aligned} \tag{89}$$

and we optimize the variational parameters in Tensorflow[9–11] using automated differentiation of the ELBO function and the BFGS algorithm.
If the covariance matrices $\boldsymbol{\Sigma}_c$ do not have hyperparameters to be optimized, i.e. $\boldsymbol{\Sigma}_c$ is fixed for all $c$, we can increase speed by pre-computing the eigendecomposition $\boldsymbol{\Sigma}_c = \mathbf{U}_c \boldsymbol{\Lambda}_c \mathbf{U}_c^{\mathrm{T}}$, such that

$$\begin{aligned}\widehat{\boldsymbol{\Sigma}}_c &= (\widehat{\sigma} N_c \boldsymbol{\Sigma}_c + \mathbf{I})^{-1} \boldsymbol{\Sigma}_c \\ &= (\widehat{\sigma} N_c \mathbf{U}_c \boldsymbol{\Lambda}_c \mathbf{U}_c^{\mathrm{T}} + \mathbf{I})^{-1} \mathbf{U}_c \boldsymbol{\Lambda}_c \mathbf{U}_c^{\mathrm{T}} \\ &= (\widehat{\sigma} N_c \mathbf{U}_c \boldsymbol{\Lambda}_c \mathbf{U}_c^{\mathrm{T}} + \mathbf{U}_c \mathbf{U}_c^{\mathrm{T}})^{-1} \mathbf{U}_c \boldsymbol{\Lambda}_c \mathbf{U}_c^{\mathrm{T}} \\ &= \left(\mathbf{U}_c (\widehat{\sigma} N_c \boldsymbol{\Lambda}_c + \mathbf{I}) \mathbf{U}_c^{\mathrm{T}}\right)^{-1} \mathbf{U}_c \boldsymbol{\Lambda}_c \mathbf{U}_c^{\mathrm{T}} \\ &= \mathbf{U}_c (\widehat{\sigma} N_c \boldsymbol{\Lambda}_c + \mathbf{I})^{-1} \boldsymbol{\Lambda}_c \mathbf{U}_c^{\mathrm{T}}\end{aligned} \tag{90}$$

such that

$$\begin{aligned}\widehat{\boldsymbol{\mu}}_c &= \widehat{\boldsymbol{\Sigma}}_c \widehat{\sigma} \overline{\mathbf{y}}_c \\ &= \widehat{\sigma} \mathbf{U}_c (\widehat{\sigma} N_c \boldsymbol{\Lambda}_c + \mathbf{I})^{-1} \boldsymbol{\Lambda}_c \mathbf{U}_c^{\mathrm{T}} \overline{\mathbf{y}}_c\end{aligned} \tag{91}$$

and the ELBO becomes

$$\mathcal{L} \propto -\frac{NG}{2}\widehat{l} - \frac{1}{2}\widehat{\sigma}\left(\sum_{g=1}^{G}\mathbf{y}_g^{\mathrm{T}}\mathbf{y}_g - 2\sum_{c=1}^{C}\widehat{\sigma}\overline{\mathbf{y}}_c^{\mathrm{T}}\mathbf{U}_c(\widehat{\sigma}N_c\mathbf{\Lambda}_c + \mathbf{I})^{-1}\mathbf{\Lambda}_c\mathbf{U}_c^{\mathrm{T}}\overline{\mathbf{y}}_c + \sum_{c=1}^{C}N_c\widehat{\sigma}^2\overline{\mathbf{y}}_c^{\mathrm{T}}\mathbf{U}_c\mathbf{\Lambda}^2(\widehat{\sigma}N_c\mathbf{\Lambda}_c + \mathbf{I})^{-2}\mathbf{U}_c^{\mathrm{T}}\overline{\mathbf{y}}_c \right.$$
$$\left. + \sum_{c=1}^{C}N_c\,\mathrm{Tr}\Big((\widehat{\sigma}N_c\mathbf{\Lambda}_c + \mathbf{I})^{-1}\mathbf{\Lambda}_c\Big)\right) - \frac{1}{2}\sum_{c=1}^{C}\Big(\widehat{\sigma}^2\overline{\mathbf{y}}_c^{\mathrm{T}}\mathbf{U}_c\mathbf{\Lambda}_c(\widehat{\sigma}N_c\mathbf{\Lambda}_c + \mathbf{I})^{-2}\mathbf{U}_c^{\mathrm{T}}\overline{\mathbf{y}}_c + \mathrm{Tr}\Big((\widehat{\sigma}N_c\mathbf{\Lambda}_c + \mathbf{I})^{-1}\Big)\Big)$$
$$-\frac{1}{2}\sum_{c=1}^{C}\log\big|(\widehat{\sigma}N_c\mathbf{\Lambda}_c + \mathbf{I})\big| + \sum_{c=1}^{C-1}N_c\widehat{v}_{2c} + \sum_{c=1}^{C}N_c\sum_{c'=1}^{c-1}\widehat{v}_{3c'} + \sum_{c=1}^{C-1}(\widehat{\alpha} - \widehat{\alpha}_{2c})\widehat{v}_{3c} - \sum_{g=1}^{G}\sum_{c=1}^{C}\widehat{\pi}_{gc}\phi_{gc}$$
$$+\sum_{g=1}^{G}\log\left(\sum_{c=1}^{C}\exp\big(\phi_{gc}\big)\right) - \widehat{\gamma}_1(1 - \log\widehat{\gamma}_2) + \log\Gamma(\widehat{\gamma}_1) + (\widehat{\gamma}_1 + 1)\widehat{l} - \sum_{c=1}^{C-1}(\widehat{\alpha}_{1c} - 1)\widehat{v}_{2c}$$
$$+\sum_{c=1}^{C-1}\log B(\widehat{\alpha}_{1c}, \widehat{\alpha}_{2c}) - (\eta_1 + C)\log\widehat{\eta}_2 + (\widehat{\eta}_2 - \eta_2)\widehat{\alpha} \tag{92}$$

### 4 Simulations and calibration/power analysis

To assess calibration of SpatialDE2's spatially variable gene detection, we first placed 625 locations on a regular grid and added a small amount of Gaussian noise to the coordinates. We then drew a size factor for each location from a uniform distribution and sampled the simulated read counts from a negative binomial distribution with a fixed dispersion and mean scaled by the size factors. This was repeated 10000 times to generate 10000 simulated non-spatially variable genes. SpatialDE2 was applied to these simulated data to obtain p-values.
To estimate statistical power of the various methods, we first trained SPARSim[12] on the mouse brain Visium data[13]. For each length scale, ranging from twice the minimal inter-location distance to the maximal inter-location distance, we drew a random pattern from a multivariate Normal distribution with a squared exponential covariance matrix. Of the 12815 genes, 1000 were defined to be spatially variable and simulated to follow the pattern. The other genes were simulated without spatial effects. Simulations were performed with SPARSim[12]. SpatialDE, SPARK, and SpatialDE2 were then applied to the simulated data set and their statistical power was estimated as the fraction of true spatially variable genes detected by the respective tool at a given FDR.

### 5 Analysis of the mouse brain and human endometrium data sets

We followed the standard ScanPy[14] workflow to filter out low-quality spots and genes. We then identified spatially variable genes on the entire tissue slice with an FDR cutoff of 0.001 and used the 2000 spatially variable genes with highest expression for segmentation. Leiden clustering was performed on the same 2000 genes after applying library size normalization, a logarithm transform to stabilize variance, principal components analysis, and nearest neighbor search as suggested in the standard ScanPy workflow. GOATOOLS[15] was used for GO enrichment analysis and false discovery rate was controlled using the Benjamini-Yekutieli procedure[16]. An FDR cutoff of 0.001 was used.
To assess the sensitivity of tissue segmentation to sequencing depth, we applied it to downsampled versions of the mouse brain data set, originally sequenced to approx. $4.5 \cdot 10^7$ reads. We compared the segmentation of the full data set to segmentation of its downsampled versions using the split/join distance[17]. The split/join distance measures the number of data points that need to be reassigned in order to obtain a consistent clustering and is more robust to subclustering than other similarity measures like the adjusted Rand index. To ensure a fair comparison, we applied each clustering method over a range of parameter values (spatial smoothness penalty for SpatialDE2 and resolution for Leiden) and selected the best setting for each sequencing depth.
For the deconvolved endometrium data, cell2location[13] output previously published in Garcia-Alonso *et al.* [18] and kindly provided by the authors was used.
