## Supplementary figures and tables for "SpatialDE2: Fast and localized variance component analysis of spatial transcriptomics"

### Supplementary Tables

**Table S1 | Features of several spatial transcriptomics analysis packages based on Gaussian Processes**

**Table S2 | Analysis of the mouse brain dataset**

**Table S3 | Analysis of the human endometrium dataset**

### Supplementary Figures

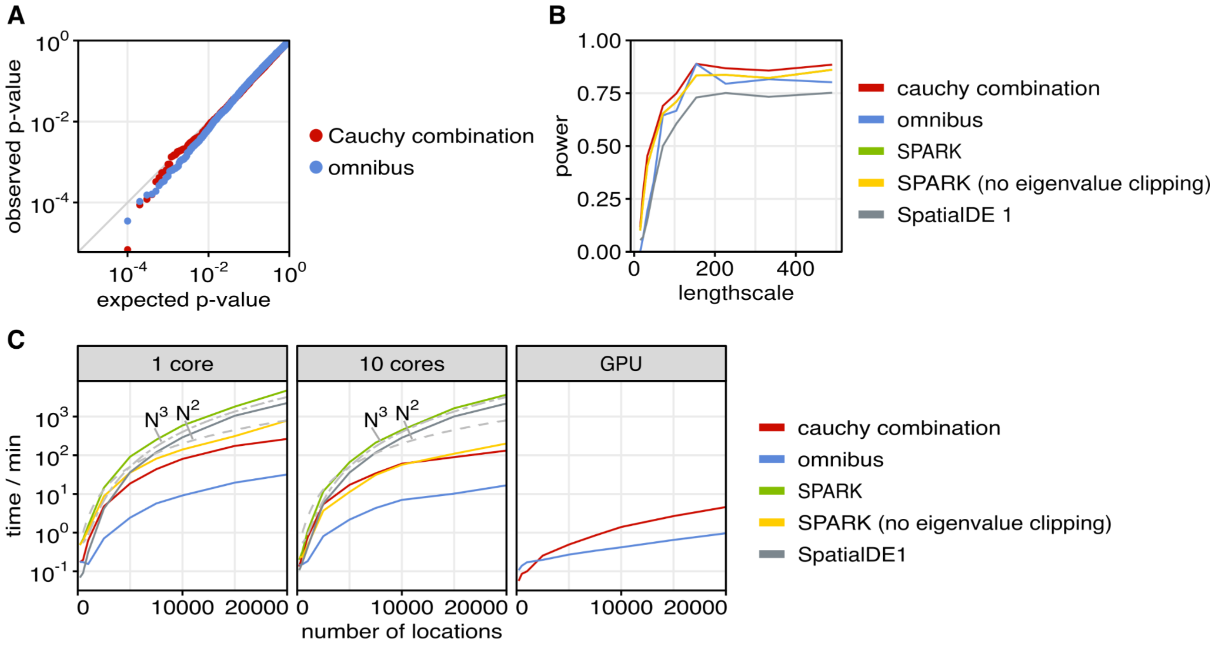


**Figure S1 | Validation and additional benchmarking results of the SpatialDE2 module for detecting spatially variable genes.** Considered are different benchmarks and assessment of spatialDE with one variance component to test for spatially variable gene expression.

(A) Q-Q plot, displaying a scatter plot between P-values expected under the null (no spatially variable expression, x-axis) versus observed P-values. Shown are results from two alternative test statistics to assess a non-zero spatial variance component in SpatialDE.

(B) Assessment of statistical power of SpatialDE2 and alternative methods for detecting spatially variable genes in simulated data with different length scales (FDR<0.05; Methods). Genes were simulated by adapting parameters from the mouse brain, simulating a four-fold maximum fold change between the highest and lowest expressed region within the simulated tissue (Methods). Considered were SpatialDE(Svensson *et al*, 2018) and SPARK(Sun *et al*, 2020), both with and without eigenvalue clipping as well as the SpatialDE2 tests. SPARK and SpatialDE tests yield comparable power.

(C) Comparison of the empirical computational complexity of SpatialDE versus Spark and SptaialDE2 using simulated data with 200 genes for increasing numbers of locations.

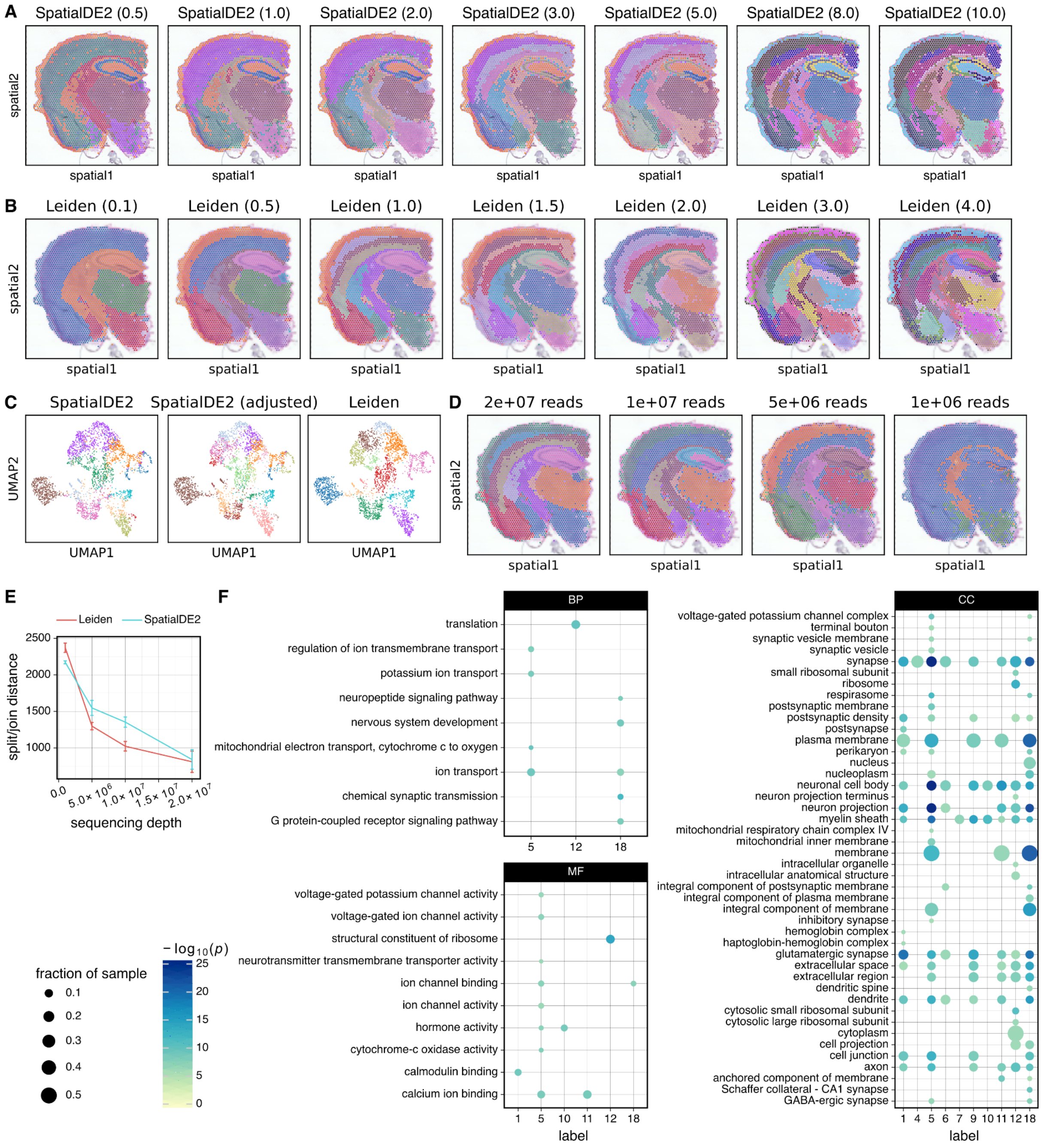


**Figure S2 | Additional results from applying SpatialDE2 to the mouse brain.**

(A,B) Effect of varying the spatial smoothness penalty on SpatialDE2 segmentation of mouse brain data. Shown are tissue region segmentations obtained using SpatialDE2 (A) and Leiden clustering (B) when varying the smoothness or resolution parameter respectively.

(C) UMAP embedding of the transcriptome profiles of individual locations of the mouse brain Visium dataset. Colour denotes tissue regions as in Fig. 3A.

(D) Results from Leiden clustering for tissue region segmentation when downsampling RNA-seq reads in the mouse brain data from (B). C.f. also main Fig. 2D for the corresponding results obtained using SpatialDE2.

(E) SpatialDE2 and Leiden clustering were applied to mouse brain visium data downsampled to the indicated sequencing depth for a range of parameter settings, and the spit/join distance compared to clustering of the full data set was calculated. For each sequencing depth, the settings yielding the lowest average distance were selected. Mean ± standard deviation of 100 resamplings is shown.

(F) GO enrichment results of region-specific spatially variable genes.

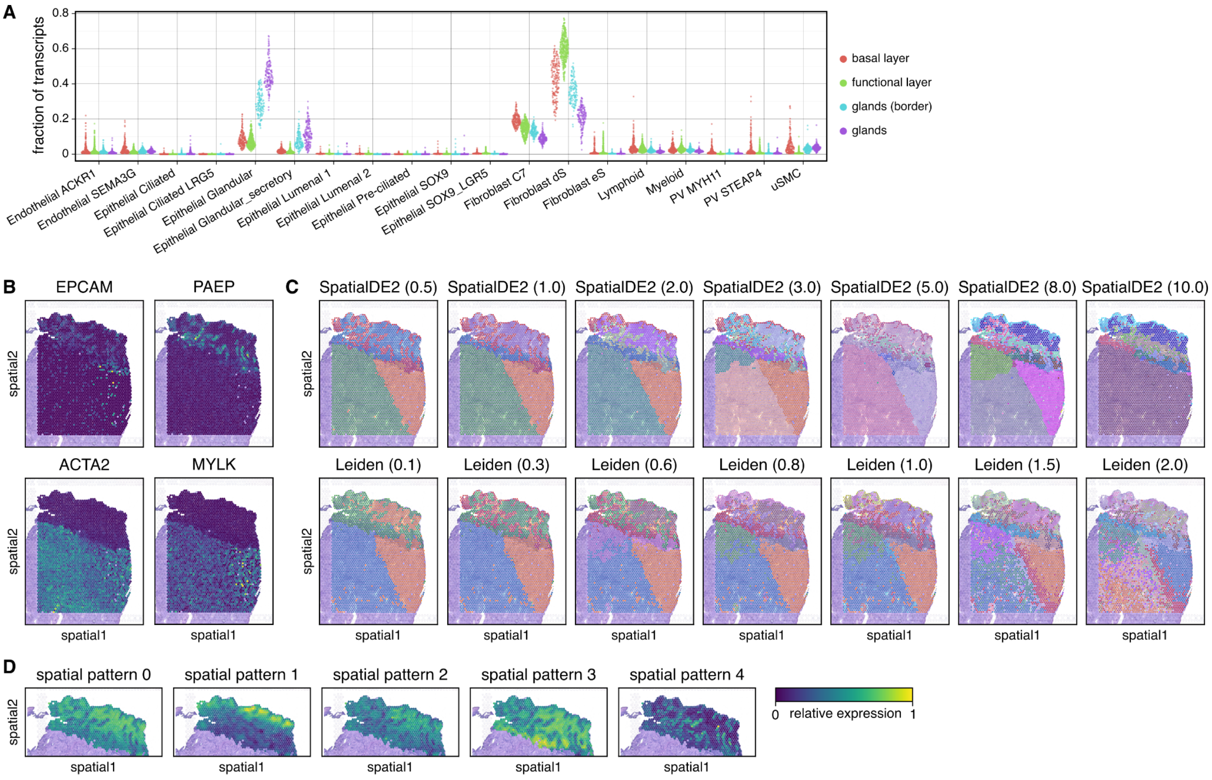


**Figure S3 | Additional results from applying SpatialDE2 to the human endometrium data.**

(A) Cell type composition of different endometrium regions identified by SpatialDE2 as estimated by cell2location.

(B) Expression of glandular cell markers (EPCAM, PAEP) and muscle cell markers (ACTA2, MYLK).

(C) Effect of varying the spatial smoothness penalty on SpatialDE2 clustering (top) and of varying the resolution parameter on Leiden clustering (bottom) of human endometrium data.

(D) Spatial expression patterns identified by SpatialDE2's automated expression histology module applied to  the basal, functional, and epithelial layers of the endometrium.

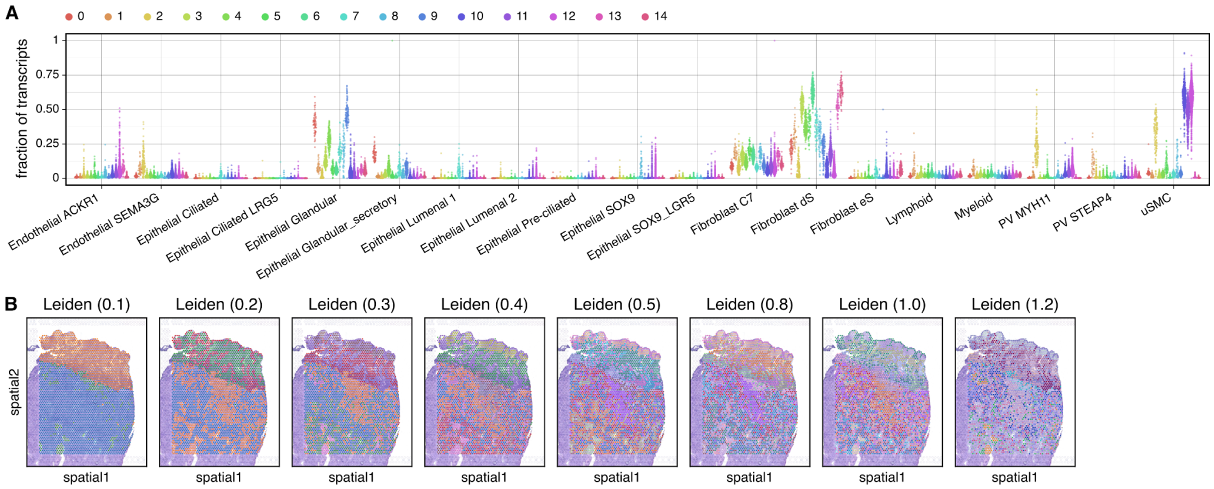


**Figure S4 | Additional results from deconvolved human endometrium data.**

(A) Cell type composition of all endometrium regions identified by SpatialDE2 as estimated by cell2location. Cf. Fig. 4A. Tissue segmentation was performed on cell2location cell type abundance estimates.

(B) Effect on varying the resolution parameter on Leiden clustering of deconvolved human endometrium data
